# RAMP-Seq: a single-tube nanopore workflow for co-amplifying full-length 16S rRNA and antimicrobial resistance genes for integrated taxonomic and resistance profiling

**DOI:** 10.64898/2026.11.12.766053

**Authors:** Deepali Pathak, Suraj Jagtap, Rekha Godbole, Nirali Patel, Ameena Mohammed, Jeevanandan Sree Priyanka, Debalina Sarkar, Imon Roy, Varsha Sridhar, Arvind Natarajan, Vishwanath Srikantaiah, Rahul Roy

## Abstract

Antimicrobial resistance (AMR) surveillance needs workflows that both identify bacteria and detect resistance genes from a single sample, but current targeted nanopore assays typically split these steps into separate reactions or library preparations. We introduce RAMP-Seq (Rapid AMR Profiling by Sequencing), a workflow that simultaneously amplifies the full-length 16S rRNA gene (∼1,500 bp) together with eight resistance determinants (∼600–700 bp) in a single multiplex PCR and then sequences the resulting mixed-length amplicon pool as a single nanopore library. The eight-primer pair set targets dominant alleles of β-lactamase, carbapenemase, quinolone-resistance and silver-resistance families. Read-length distributions confirmed that both size classes were co-amplified evenly using a single, non-equimolar primer pool. From extracted DNA to the first taxonomically classified reads took 8.2 ± 0.6 h. Full-length 16S sequencing of an eight-member mock community identified all anticipated taxa, and the observed abundances closely followed the expected log-scaled composition (Pearson’s r = 0.879). The full nineplex amplicon assay was subsequently tested on three distinct matrices: pooled clinical isolates, spiked human blood, and environmental water samples. Using an equal-mass pool of genomic DNA from eleven multidrug-resistant clinical and repository isolates, the full assay produced genus-level and resistance-gene profiles that were consistent across biological replicates (*r* = 0.998 and *r* = 0.824), and it specifically identified all seven panel targets present, confirmed by Illumina whole-genome sequencing. After carbonate-based host DNA depletion, *Klebsiella pneumoniae* was recovered from spiked human blood at every titration down to 10² CFU/mL, as were all five panel targets carried by that strain. Across five water-sampling sites from a sewage treatment plant (STP) to a connected lake system, microbial communities and resistance-gene profiles were site-specific: influent and effluent communities overlapped little, while the lake inlet and outlet were similar. Target resistance genes were most numerous and abundant in STP influent, fewer in effluent, and rare or absent in lake and well samples. Because bacterial identification relies on amplification of the conserved, near-universal 16S rRNA gene, rather than on species-specific primer sets, RAMP-Seq can detect and taxonomically classify bacteria in a design-agnostic manner. The same single-tube chemistry therefore serves sample types as different as clinical specimens and environmental water and is extendable to longer ribosomal targets and further resistance-gene families.

**Impact statement:** Antimicrobial resistance surveillance must simultaneously identify bacteria and their resistance genes, but most sequencing workflows require separate reactions. This limits throughput and increases costs, constraints especially relevant for low- and middle-income settings. RAMP-Seq combines both tasks in one multiplex PCR, amplifying full-length 16S rRNA for identification plus eight resistance-gene targets, then sequencing them together as a single nanopore library on a portable MinION. From extracted DNA to first classified reads takes ∼8 hours, and one barcode per sample means a 12-barcode kit supports 12 samples. We successfully applied RAMP-Seq on multidrug-resistant clinical isolates, bacteria spiked into human blood, and wastewater and freshwater samples. Because it relies on a universal bacterial marker, it is not limited to predefined species, enabling use when pathogens are unknown, and lab capacity is limited.

**Data Summary:** All sequencing datasets produced in this study have been deposited in the NCBI Sequence Read Archive (SRA) under BioProject accession PRJNA1483527. Oxford Nanopore reads include the multidrug-resistant clinical isolate pool (two biological replicates; SRR39373345–SRR39373346); environmental water collected from five locations across a sewage treatment plant and the connected lake system (14 runs including technical replicates; SRR40992843–SRR40992856); and *Klebsiella pneumoniae* spiked into human blood at 10⁶–10¹ CFU/mL (two biological replicates per dilution; accessions SAMN64748836–SAMN64748837). **Supplementary Tables S1–S10** contain culture-based identification and antimicrobial susceptibility results for the clinical isolates (**S1**), environmental site descriptions and coordinates (**S2**), sequences of the panel primers and of primers excluded during development (**S3–S4**), primer concentrations (**S5**), *in silico* coverage of the primer panel (**S6**), summaries of sequencing runs and replicates (**S7)**, Illumina whole-genome sequencing results (**S8**), read-length binning statistics (**S9**), and genus-level profiles of environmental samples (**S10**). All data, materials, and protocols required to reproduce the results are available in the article, the supplementary materials, and the repositories listed above.

## Introduction

Bacterial antimicrobial resistance (AMR) is among the leading causes of death worldwide [1]. The latest Global Burden of Disease analysis projects 8.22 million associated deaths and 1.91 million attributable deaths by 2050, with south Asia among the super-regions expected to experience the highest mortality rates [2]. Carbapenem resistance in Gram-negative bacteria has increased more rapidly than resistance to any other antibiotic class, with an estimated 1.03 million related deaths in 2021. [2]. Comprehensive population-based surveillance is nonetheless scarce where the burden is greatest across low- and middle-income countries where weak laboratory capacity, limited health information systems, and constrained resources leave the available data fragmented and unrepresentative [3]. Resistance is not limited only to clinical settings. The same drug resistance determinants circulate between human, animal and environmental reservoirs on mobile genetic elements, moving between wastewater discharge, agricultural runoff and direct contact [4, 5], and urban wastewater has been used to monitor the abundance of resistance genes across dozens of countries [6]. This interconnection gives One Health real implications for surveillance: a resistance gene without its host organism, or a pathogen without its resistance genes, is only half the picture especially if it cannot be traced across compartments. However, key compartments can differ dramatically, eg. blood is dominated by host DNA, while surface water is diverse and low-biomass. One Health surveillance therefore needs methods that can jointly report bacterial identity and resistance genes across dissimilar matrices, enabling direct comparison of clinical and environmental results.

Current molecular approaches to AMR detection address only part of this challenge. Culture-based antimicrobial susceptibility testing remains the clinical reference standard but is slow, limited to cultivable organisms, and unable to resolve the genetic basis of resistance [7]. Whole-genome sequencing (WGS) provides comprehensive resistome characterisation but requires substantial laboratory infrastructure, bioinformatic expertise, and turnaround times that are incompatible with rapid clinical decision-making [8]. In clinical settings, it is usually carried out on a cultured isolate, and therefore shares the same delay and the need for the organism to be cultivable as the preceding step. PCR-based assays offer speed but typically target narrow gene sets, sacrificing breadth for sensitivity. On the other hand, shotgun metagenomics captures both taxonomic and resistance information. However, its high per-sample cost, sequencing depth requirements, and computational demands limit scalability, particularly in resource-constrained settings [8]. Targeted amplicon sequencing sits between these two extremes, retaining the sensitivity and low input requirements of PCR while providing sequence-level readouts and enabling multiple targets to be analysed in a single assay. For environmental surveillance, high-throughput qPCR arrays allow quantitative and parallel resistance-gene profiling [9, 10], but they report only the targets included on the array and usually do not provide any taxonomic context. Although nucleic-acid sequencing capacity grew markedly during the COVID-19 pandemic, yet remains unevenly distributed [11], and workflow and protocols built from multiple reaction steps, separate library preparations, and specialist handling keep per-sample cost and complexity high enough to restrict routine use in pathogen surveillance.

Targeted nanopore sequencing has emerged as a practical approach for rapid, field-ready AMR detection, and multiple workflows have been reported [12–14]. MultiSeq-AMR is the most comprehensive, covering 91 resistance gene groups across 11 antibiotic classes and validated on clinical bloodstream samples, but it uses 13 primer pools plus a 14th for species markers, requiring multiple reactions and barcodes per sample [13]. TAROT applies core-genome SNP-based amplicon tiling to nosocomial transmission tracking and does not attempt broad taxonomic profiling [12], whereas RapidONT streamlines clinical whole-genome sequencing but retains the infrastructure requirements and turnaround of a genome-scale approach [14]. For taxonomic identification and resistance gene detection, sequencing samples are generated in separate reactions with separate primer pools or as separate libraries, so the number of reactions and barcodes consumed scales with the breadth of the panel rather than with the number of samples. To our knowledge, no reported workflow co-amplifies a full-length taxonomic marker together with short resistance-gene amplicons in a single reaction and sequences them as one library.

We present RAMP-Seq (Rapid AMR Profiling by Sequencing), which co-amplifies the full-length 16S rRNA gene (∼1,500 bp) and eight short resistance targets (∼600–700 bp) in a single multiplex PCR and sequences them as one nanopore library, requiring one reaction and one barcode per sample. The key technical challenge is to ensure uniform recovery across this fragment-size range, because shorter products would otherwise be preferentially amplified. In RAMP-Seq, this is achieved using a single, non-equimolar primer pool in which the full-length 16S primer pair is kept at the highest concentration, along with cycling conditions optimised for the longest amplicon. The panel comprises **t**he CTX-M-1-group extended-spectrum β-lactamase (*bla_CTX-M-1_*), class A β-lactamases (*bla*_TEM_, *bla*_SHV/LEN/OKP_), carbapenemases (*bla*_NDM-1_, *bla*_OXA-48_), an AmpC β-lactamase (*bla*_CMY-6_), a plasmid-mediated quinolone-resistance determinant (*qnrB1*) and a silver-resistance determinant (*silP*), selected for their clinical and environmental prevalence in South Asia [15–18]. As our primers bind conserved regions, their predicted coverage extends to over 499 related resistance genes. We tested the workflow using a defined mock community, a set of ten multidrug-resistant clinical isolates, *Klebsiella pneumoniae* spiked into human blood, and both wastewater and freshwater samples, evaluating taxonomic accuracy, agreement of resistance-gene calls with whole-genome sequencing, analytical sensitivity in a host-rich matrix, and performance across different sample types.

## Methods

### Study Design and Sample Sources

RAMP-Seq performance was evaluated using four primary validation datasets: (i) a ZymoBIOMICS mock microbial community for assessment of taxonomic accuracy, (ii) an equal-mass pool of multidrug-resistant clinical isolates for integrated taxonomic and AMR profiling, (iii) *Klebsiella pneumoniae* spiked into human blood for analytical sensitivity testing, and (iv) environmental water samples spanning a wastewater treatment-to-freshwater gradient collected from a sewage treatment plant and downstream lake system in Bengaluru, India.

Separately, exploratory experiments using *Escherichia coli* and amplification of the longer 16S-ITS-23S rRNA operon were performed to assess compatibility of the workflow with longer ribosomal targets and were not included in formal performance evaluation.

MDR clinical isolates were obtained from the Department of Clinical Microbiology, Sri Devaraj Urs Academy of Higher Education and Research (SDUAHER), Kolar, India. Culture-based identification and antimicrobial resistance profiles are shown in **Supplementary Table S1**. Each isolate was identified by its laboratory specimen number followed by a suffix (B1, B2) for organisms recovered from the same specimen; for example, BC/1553/2021-B1 (*Escherichia coli*) and BC/1553/2021-B2 (*Enterococcus* spp.) were both isolated from one blood culture. Environmental samples, including sewage treatment plant influent and effluent, lake inlet and outlet, and open well water, were collected across Bengaluru by Molecular Solutions Care Health LLP. Sampling locations and metadata are provided in **Supplementary Table S2**. DNA extracted from all sample types were transported on ice to the Indian Institute of Science (IISc), Bengaluru, and stored at −80°C until processing.

All experiments other than the mock community were performed with the complete nine-target reaction so that taxonomic and resistance profiling were generated together from a single amplicon library. The ZymoBIOMICS mock community standard does not contain panel resistance determinants, so the AMR primers were omitted, and this dataset was used to standardise full-length 16S amplification and classification. The clinical isolate pool, which carried panel targets, therefore provided the integrated evaluation of the complete assay, while the blood-spike and environmental experiments additionally assessed analytical sensitivity in a host-rich matrix and performance across sample types.

All experiments were performed with a minimum of two independent biological replicates. Biological replicates (BR) represent independent sample preparations originating from separate DNA extractions of the same sample type, followed by independent multiplex PCR, library preparation, and sequencing runs. Technical replicates (TR) represent independent multiplex PCR amplifications and sequencing runs generated from the same DNA preparation. Unless otherwise stated, statistical analyses were performed using biological replicates, whereas technical replicates were included solely to assess workflow reproducibility. Technical replicate sequencing runs were performed for selected wastewater samples during assay optimisation; these data were not treated as independent biological observations during statistical analyses.

Environmental samples were collected on 30 August 2023 from five sites spanning a wastewater-to-freshwater gradient: Raja Canal sewage treatment plant influent (RC_In_) and effluent (RC_Out_), Bagalur Lake inlet (BL_In_) and outlet (BL_Out_), and a nearby open well (BL_OW_) (**Supplementary Table S2**). These samples were used to evaluate whether the workflow works effectively across environmental matrices that vary in turbidity, biomass, and community complexity. The replicates are independent extractions and amplifications from each sample rather than repeated sampling events, so the results describe assay performance rather than a measurement of environmental resistome.

### Bacterial Strains and Control

Strains were selected to meet two requirements of assay development: coverage of the organisms that carry the greatest burden of antimicrobial resistance and carriage of the resistance determinants included in the panel. The selected taxa—*Escherichia coli, Klebsiella pneumoniae, Acinetobacter spp., Pseudomonas aeruginosa, Enterococcus spp. and methicillin-resistant Staphylococcus aureus*—span Gram-negative and Gram-positive organisms and correspond to pathogens designated critical or high priority in the WHO bacterial priority pathogens list [19], five of which also account for the majority of deaths attributable to bacterial AMR [1].

For multiplex 16S rRNA and AMR PCR validation, genomic DNA from multidrug-resistant clinical isolates was used, including representatives of *Escherichia coli, Klebsiella pneumoniae, Acinetobacter spp., Pseudomonas aeruginosa, Enterococcus* spp., and multidrug-resistant *Staphylococcus aureus* (**Supplementary Table S1**). These isolates were selected for their relevance to bloodstream infections and their known carriage of target resistance genes included in the RAMP-Seq panel. The reference strain *Klebsiella pneumoniae* MCC 2451 (= ATCC 700603, strain K6; originally isolated from the urine of a hospitalised patient, USA), procured from the National Centre for Microbial Resource (NCMR), NCCS, Pune, was used as a preliminary control for primer standardisation and validation.

Spike-in experiments to assess the analytical sensitivity of 16S rRNA were performed primarily using *Klebsiella pneumoniae* (ET/167/2021-B1, source: endotracheal aspirate). Defined bacterial loads 10^6^ −10^1^ CFU/mL) were prepared by serial dilution and introduced into human blood to simulate low-biomass clinical samples.

To evaluate long amplicon compatibility, including amplification of the 16S–ITS–23S rRNA operon, genomic DNA from representative *Escherichia coli* (BC/1553/2021-B1) was used in exploratory single-tube PCR assays. In addition, spike-in experiments targeting the full-length 16S–ITS–23S rRNA operon region were performed using *Escherichia coli* (10^6^ −10^1^ CFU/mL), enabling assessment of assay performance for longer ribosomal amplicons under low-biomass conditions.

### Ethical Approvals

All clinical isolates were anonymised before transfer. Ethical clearance for the use of clinical isolates was granted by the SDUAHER Institutional Ethics Committee (Approval No. IEC/SDUAHER/2021/07). Environmental sampling was carried out under authorisation from the Bangalore Water Supply and Sewerage Board (BWSSB; Ref. No. BWSSB/CE(P)/ACE(P)-1/TA/Q112/202324). All laboratory procedures were conducted in accordance with the Institutional Biosafety Committee at IISc (Approval No. IBSC/IISc/RR/40/202324). Human blood used for spiking experiments was obtained and used under Institutional Ethics Committee Approval No. 29 for a study on multiplex PCR-based detection of drug resistance.

### DNA extraction

Genomic DNA from multidrug-resistant (MDR) clinical isolates was extracted using the QIAamp DNA Host-Free Microbiome Kit (Qiagen, Cat. No. 51904), suitable for polymicrobial and low-biomass samples. DNA from blood spike-in experiments was isolated using the DNeasy Blood & Tissue Kit (Qiagen, Cat. No. 69504), while plasmid-enriched preparations were obtained using the QIAprep Spin Miniprep Kit (Qiagen, Cat. No. 27106X4), as appropriate for the sample type. Genomic DNA from the reference strain *Klebsiella pneumoniae* MCC 2451 (= ATCC 700603; NCMR, NCCS, Pune) was extracted with the QIAamp DNA Host-Free Microbiome Kit and served as a positive control.

Environmental samples were centrifuged at 10,000×g for 15 min at room temperature to pellet biomass. The resulting pellets were carefully retained, and genomic DNA was extracted using the FastDNA SPIN Kit for Soil (Cat. No. 116560200, MP Biomedicals) according to the manufacturer’s instructions.

DNA concentration was measured using a Qubit 4 Fluorometer (Thermo Fisher Scientific) with the Qubit dsDNA High Sensitivity Assay (Thermo Fisher Scientific, Cat. No. Q33231). DNA purity was assessed by measuring the A260/280 absorbance ratio using a NanoDrop spectrophotometer (Thermo Fisher Scientific). DNA integrity and successful recovery of bacterial DNA were further confirmed by agarose gel electrophoresis and PCR amplification of the 16S rRNA gene.

### Primer Design and Target Selection

Initially, twenty-three primer pairs were designed to target major antimicrobial resistance gene families, including extended-spectrum β-lactamases (*bla_CTX-M-1_, bla_TEM_, bla_SHV/LEN/OKP_*), carbapenemases (*bla_NDM-1_, bla_OXA-48_*_)_, AmpC β-lactamase (*bla_CMY-6_*), plasmid-mediated quinolone resistance (*qnrB1*_)_, and a silver-resistance determinant (*silP*), alongside universal full-length 16S rRNA gene primers (27F/1492R). Primers were designed using Primer3 [20] and NGS-PrimerPlex [21], and specificity was verified using NCBI Primer BLAST [22].

Amplification specificity was confirmed by agarose gel electrophoresis (single band of expected size, absence of non-specific products) and Sanger sequencing. Of the twenty-three candidates, eight AMR targets that consistently produced strong, specific amplification without detectable cross-reactivity were selected for the final multiplex panel **(Supplementary Figure S1)**. The remaining fifteen primer pairs were excluded due to weak amplification, non-specific products, or primer-dimer formation under multiplex conditions. All primer pairs evaluated during assay development, including those excluded due to non-specific amplification or poor multiplex performance, are listed in **Supplementary Tables S3** and **S4**. Final primer concentrations are given in **Supplementary Table S5**, and the predicted *in silico* coverage of the panel primers is given in **Supplementary Table S6**.

### Multiplex PCR Co-amplification

Multiplex PCR was performed to co-amplify the full-length 16S rRNA gene and eight targeted antimicrobial resistance (AMR) determinants in 50 µL reactions containing 1× LongAmp Taq 2X Master Mix (New England Biolabs, Cat. No. M0287S), 50 ng of template DNA, and a single non-equimolar primer pool in which the eight AMR primer pairs were present at final concentrations of 0.4-0.8 µM each and the full-length 16S rRNA primer pair at 2.0 µM **(Supplementary Table S5**). This primer configuration was selected to enable recovery of the ∼1,500 bp 16S rRNA amplicon alongside the shorter AMR amplicons. LongAmp Taq was selected for its robust processivity across the ∼607-1,506 bp amplicon size range, enabling simultaneous amplification of short AMR targets and the full-length 16S rRNA gene within a single reaction. Final primer concentrations for each target are listed in **Supplementary Table S5**. Thermocycling for 16S rRNA-AMR co-amplification consisted of an initial denaturation at 94°C for 30 s, followed by 25 cycles of 94°C for 60 s, 60°C for 60 s, and 65°C for 60 s, with a final extension at 65°C for 10 min.

For the exploratory 16S-ITS-23S rRNA operon experiments, co-amplification of the 16S-ITS-23S region and AMR targets was performed in 50 µL reactions containing 50 ng of template DNA and 1× LongAmp Taq 2× Master Mix. The thermal cycling profile consisted of an initial denaturation at 98°C for 30 s, followed by 25 cycles of denaturation at 98°C for 7 s, annealing at 59°C for 30 s, and extension at 72°C for 150 s, with a final extension at 72°C for 10 min.

PCR products were visualised by electrophoresis on 1.5% agarose gels alongside a 100 bp DNA Ladder (New England Biolabs, Cat. No. N3231S) or, for the 16S-ITS-23S rRNA operon products, a 1 kb DNA Ladder (New England Biolabs, Cat. No. N3232S) and matched the expected amplicon sizes (Supplementary Table S3). Amplification balance between long (16S rRNA and 16S-ITS-23S rRNA operon) and short (AMR) targets was assessed using gel band intensities and post-sequencing read-length distributions (**Figures 1d–e** and **Supplementary Figures S2a** and **S2b)**.

**Figure 1:**
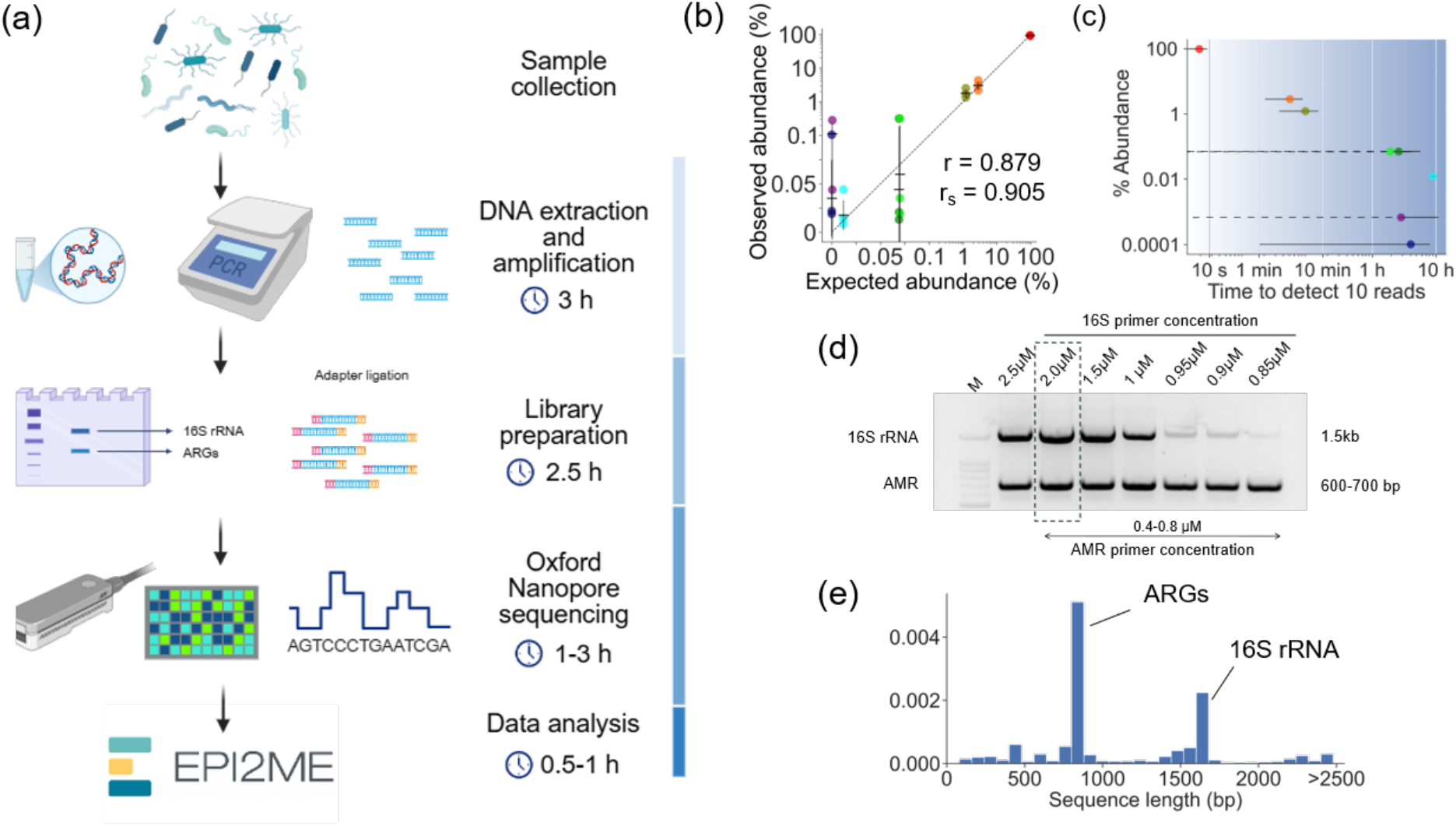
Establishment of Rapid AMR Profiling by Sequencing (RAMP-Seq). **(a)** Overview of the RAMP-Seq workflow: sample collection, DNA extraction and multiplex PCR, library preparation, Oxford Nanopore sequencing, and data analysis. The approximate time required for each step is indicated. **(b)** Observed versus expected relative abundances in an eight-member mock microbial community with a logarithmic abundance distribution, amplified with the universal 27F/1492R primers alone (Pearson’s r = 0.879; Spearman’s r^s^ = 0.905). Error bars represent the standard deviation of two biological replicates. **(c)** Time required to obtain the first 10 taxonomically assigned reads for each organism in the mock community as a function of relative abundance. Error bars represent the standard deviation; dashed lines indicate values crossing zero. **(d)** Multiplex PCR products generated with 16S rRNA primer concentrations of 0.85–2.5 µM and AMR primer concentrations of 0.4–0.8 µM, used to identify conditions giving balanced amplification of the 16S rRNA gene (∼1.5 kb) and the eight AMR targets (600– 700 bp). The dashed box marks the 2.0 µM 16S primer concentration used in the final non-equimolar primer pool. Target AMR genes included *bla_SHV/LEN/OKP_*, *bla_CMY-6_*, *bla_NDM-1_*, *bla_CTX-M-1_*, *bla_TEM_*, *qnrB1*, *bla_OXA-48_*, and *silP;* expected amplicon sizes are listed in **Supplementary Table S3a**. M, 100 bp DNA ladder. **(e)** Read-length distribution of the multiplex reaction, showing two dominant peaks corresponding to the short AMR amplicons and the full-length 16S rRNA amplicon.

### Nanopore Library Preparation and Sequencing

PCR products were purified with the QIAquick Nucleotide Removal Kit (Qiagen, Cat. No. 28304) and quantified on a Qubit 4 Fluorometer (Thermo Fisher Scientific). Individual samples were barcoded through the ONT Universal Tag on the panel primers **(Supplementary Table S3b)** in a second PCR using the PCR Barcoding Expansion Kit (Oxford Nanopore Technologies, EXP-PBC001) and LongAmp Taq 2X Master Mix (New England Biolabs, Cat. No. M0287S) (95°C for 3 min; 15 cycles of 95°C for 15 s, 62°C for 15 s and 65°C for 5 min; final extension at 65°C for 5 min), and the barcoded products were cleaned up with AMPure XP beads (Beckman Coulter, Cat. No. A63881). Libraries were prepared with the Oxford Nanopore Technologies (ONT) Ligation Sequencing Kit (SQK-LSK114). Barcoded amplicons were pooled to a total input of 20–50 fmol and sequenced on MinION R10.4.1 (FLO-MIN114) or Flongle R10.4.1 (FLO-FLG114) adapters (sample-to-platform assignments are specified in **Supplementary Table S7**). Sequencing was performed on the MinION Mk1B device, and FASTQ files were generated in real time using MinKNOW v23.07 with integrated live basecalling. Run summaries, including total reads per sample and their biological replicates, are provided in **Supplementary Table S7**.

### Mock Community Validation

Mock-community validation was performed using the ZymoBIOMICS Microbial Community DNA Standard II (Log Distribution; Zymo Research, Cat. No. D6311) to assess taxonomic profiling accuracy. Full-length 16S rRNA amplicons were therefore generated from 50 ng of template DNA using ONT adapter-tailed 27F/1492R primers and processed through the remainder of the RAMP-Seq workflow (library preparation, sequencing, read filtering and classification) under standard conditions (**Supplementary Table S7)**. The observed taxonomic profiles were compared with the manufacturer’s expected composition.

### Illumina Sequencing

Whole-genome sequencing (WGS) was performed on seven of the clinical isolates using the Illumina NovaSeq 6000 platform (RRID: SCR_016387), generating paired-end reads (2×150 bp). Reads were trimmed based on quality (Q20+) using Geneious Prime (v2024.0.1) and mapped to the antimicrobial-resistant gene database MEGARes v2.0 [23] using Geneious aligner with Medium/Low sensitivity. Genes with a minimum average depth of 5X and coverage breadth of 80% are reported in **Supplementary Table S8**. Agreement between RAMP-Seq and Illumina WGS was assessed at the qualitative target level (detected/not detected) for the eight AMR determinants included in the multiplex panel. Comparisons were performed on a pooled clinical-isolate dataset rather than on an isolate-by-isolate basis. A target was considered concordant when the corresponding determinant detected by RAMP-Seq was also identified in the Illumina WGS dataset of the clinical isolates containing that determinant.

A preliminary taxonomy assignment was performed based on Kraken2 [24] with Standard-8 dataset (**Supplementary Table S8a**). The reference sequence from the identified taxa were used to map the Illumina reads using Geneious aligner with medium/low sensitivity (**Supplementary Table S8**). The Illumina reads were also mapped to 16S rRNA gene sequences of the Kraken identified taxa (**Supplementary Table S8b**).

### Spiked Blood Titration

To evaluate analytical sensitivity in a clinically relevant matrix, *Klebsiella pneumoniae* ET/167/2021-B1 was cultured to late log phase in Luria–Bertani (LB) broth (HiMedia, Cat. No. M1245) at 37°C with shaking, following standard bacterial growth protocols. Bacterial density was estimated by measuring the optical density at 600 nm (OD₆₀₀). The culture was then serially diluted in sterile PBS, and aliquots corresponding to target concentrations of 10^6^ −10^1^ CFU/mL were plated and retrospectively quantified by colony counting.

Based on the confirmed viable cell counts, a bacterial suspension containing the corresponding bacterial concentration was prepared and spiked into 1 mL of EDTA-treated human blood to establish the starting inoculum. This spiked blood sample was then serially diluted tenfold directly in blood to obtain a dilution series ranging from 10^6^ −10^1^ CFU/mL. Final bacterial concentrations used for spike-in experiments were confirmed retrospectively by colony-forming unit (CFU) enumeration following overnight culture of plated dilutions.

To simulate real-world sample handling conditions prior to laboratory processing, spiked blood samples were incubated at room temperature for 30 min before downstream processing. All spiking and sample-handling steps were performed under sterile conditions in a laminar flow hood. Unspiked blood served as a negative control.

In the analytical sensitivity experiments, the lowest concentration detected in both biological replicates was considered the lowest reproducibly detectable concentration under the tested conditions. This value is not a formally validated limit of detection, but the lowest concentration reproducibly detected in both replicates.

### Host DNA Depletion

Host DNA was depleted using a rapid carbonate-based lysis protocol [25]. Briefly, samples were treated with a carbonate lysis buffer containing 2M Na₂CO₃ (Sigma-Aldrich, Cat. No. 223465) and 1% Triton X-100 (Sigma-Aldrich, Cat. No. T8787) for 3 min at room temperature, followed by neutralisation with 1M Tris-HCl (pH 4.5). Samples were then centrifuged at 5,000×g for 5 min, and the resulting bacterial pellets were resuspended in PBS (Gibco, Cat. No. 10010023). DNA was extracted using the QIAamp DNA Blood Mini Kit (Qiagen, Cat. No. 51104). Two biological replicates were processed at each dilution. Host DNA depletion efficiency and bacterial DNA recovery were assessed by agarose gel electrophoresis of 16S rRNA PCR products from *Escherichia coli-*spiked blood (**Supplementary Figure S3**), Qubit-based DNA quantification, and PCR amplification of the 16S rRNA gene.

### Nanopore Data Analysis

Basecalling was performed in real time using the *dna_r10.4.1_e8.2_400bps_hac* model in MinKNOW v23.07. Base-called reads with a minimum quality score of Q9 were retained for analysis. This permissive threshold was chosen to maximize sensitivity for the short AMR amplicons, which require lower per-read accuracy when alignment-based detection is performed against a curated reference database. All analyses were performed using the EPI2ME Metagenomics Workflow v2.10.1 (Oxford Nanopore Technologies; RRID: SCR_016382; https://epi2me.nanoporetech.com/).

For timing analyses, the first taxonomic assignment was defined as the earliest timepoint at which sequencing reads satisfying the predefined quality and classification criteria were assigned to a bacterial taxon by the EPI2ME Metagenomics Workflow [26]. Workflow duration was measured from introduction of extracted DNA into the multiplex PCR reaction to generation of the first taxonomically classified sequencing reads following sequencing, basecalling, and automated analysis.

Reads were separated by length into two categories: long reads (1,250-1,750 bp), corresponding to full-length 16S rRNA amplicons, and short reads (500-1,000 bp), corresponding to AMR amplicons. Length-based filtering was performed using EPI2ME. Reads falling outside these windows were excluded from downstream analysis. These windows were selected to encompass the expected sizes of the AMR amplicons (600-800 bp) and full-length 16S rRNA amplicons (∼1,500 bp) while excluding primer-dimers, truncated amplification products, and non-target amplicons. The excluded fraction varied between samples (**Supplementary Table S9**). For the exploratory 16S-ITS-23S experiments, reads were separated using a 3,000-5,000 bp window corresponding to the expected ∼4.2-4.5 kb operon amplicon.

### Taxonomic classification

Long reads were aligned to the NCBI 16S rRNA reference [27, 28] using Minimap2 v2.24 [29] in the EPI2ME Metagenomics Workflow v2.10.1 [26]. Relative abundance was calculated as the proportion of classified reads assigned to each taxon within each sample. Taxa with >1% reads mapping and >5 reads mapping were reported. All other detected taxa were grouped as “other”. Reads unmapped to any 16S rRNA reference sequence were excluded from taxonomic assignment and labelled as unclassified. The 1% abundance threshold was selected based on pooled bacterial samples to minimise spurious taxonomic assignments while retaining sufficient sensitivity for genus-level classification from Nanopore full-length 16S rRNA reads.

### AMR detection

Short reads were aligned to the MEGARes v2.0 database [23] using Minimap2 v2.24 [29] in the EPI2ME Metagenomics Workflow v2.10.1 with an identity threshold of 80% and a minimum query coverage threshold of 50%. AMR relative abundance was calculated as the number of reads assigned to a given resistance determinant divided by the total number of short reads mapped to any AMR target within that sample. Relative abundances therefore represent within-sample proportions of targeted AMR-associated reads and should not be interpreted as measures of absolute gene abundance.

The alignment thresholds were selected to balance sensitivity and specificity for Nanopore-derived amplicons while accommodating the residual per-read error profile characteristic of long-read sequencing error.

Unspiked blood controls and no-template controls (NTCs) were included in all sequencing runs. For blood-spike experiments, one unspiked blood control was processed alongside each biological replicate series, and at least one NTC was included in each multiplex PCR run. A target was considered detected when at least one read satisfying the predefined alignment thresholds was assigned to the corresponding taxon or resistance determinant. A target was considered absent when no reads meeting these criteria were detected. No, taxonomically classified long reads or AMR-associated short reads meeting the detection criteria were identified in any unspiked blood control or no-template control.

### Statistical Analysis

Pearson correlation coefficients (r) and Spearman’s rank correlation coefficients (r_s_) were used to assess reproducibility across biological replicates and concordance between observed and expected compositions. Where applicable, results are reported as mean ± standard deviation from independent biological replicates. Statistical analyses were performed in Python v3.12 using SciPy Stats.

## Results

### Establishment of RAMP-Seq

RAMP-Seq co-amplifies taxonomic and resistance targets in a single multiplex PCR so that one reaction and one barcode per sample yield both profiles from the same nanopore library (**Figure 1a**). Across multiple independent runs, the complete RAMP-Seq workflow from extracted DNA to generation of taxonomically classified sequencing reads was completed in 8.2 ± 0.6 hours (**Figure 1a**).

We first established the taxonomic component of the assay. Universal 27F/1492R primers amplified the full-length 16S rRNA gene as a single product of the expected ∼1,500 bp (**Supplementary Figure S1**). Because longer ribosomal targets offer finer taxonomic resolution, we also amplified the complete 16S–ITS–23S rRNA operon from *Escherichia coli* genomic DNA, which yielded a clean band of ∼4.2–4.5 kb (**Supplementary Figure S2**), showing the reaction chemistry to be compatible with ribosomal amplicons of this length.

Taxonomic accuracy was then benchmarked against the ZymoBIOMICS mock community standard, using the universal 27F/1492R primers alone. Since the standard carries no acquired resistance genes, these runs were to report the reliability of taxonomic components by amplicon sequencing. All eight expected taxa were correctly identified, with strong agreement between observed and expected relative abundances (Pearson’s *r* = 0.879, Spearman’s *r_s_* = 0.905; **Figure 1b**). Sequencing the 16S–ITS–23S operon from the community recovered 6 of the 8 species, with the abundances of those six closely matching expectations (Pearson’s *r* = 0.914, Spearman’s *r_s_* = 0.986; **Supplementary Figure S4**). Taxa present at >1% relative abundance were detected within 10 ± 2 minutes of sequencing, defined as the time required to obtain the first 10 taxonomically assigned reads for a given organism, whereas low-abundance taxa required up to 9.4 ± 0.8 hours to accumulate sufficient classified reads (**Figure 1c**). Both values are based on two independent biological replicates and are measured from library loading, not from receipt of the sample.

We next assembled the resistance component. Twenty-three primer pairs targeting antimicrobial resistance determinants were screened individually, of which eight AMR (*bla_SHV/LEN/OKP_*, *bla_CMY-6_*, *bla_NDM-1_*, *bla_CTX-M-1_*, *bla_TEM_*, *qnrB1*, *bla_OXA-48_*, and *silP*) consistently yielded specific and reproducible amplification and were therefore included in the final multiplex panel (**Supplementary Figure S1**, **Supplementary Table S3a**). Amplicon sizes (607–1,506 bp) were confirmed by gel electrophoresis and Sanger sequencing. The remaining primer pairs were excluded because they failed to generate reliable amplification during assay development or produced amplification under singleplex conditions but could not be reproducibly detected following incorporation into the multiplex assay. Several corresponding resistance genes were subsequently identified by Illumina whole-genome sequencing of the clinical isolates, indicating that limitations in primer compatibility, target-sequence diversity, or amplification efficiency, rather than the absence of the underlying resistance determinants likely contributed to their exclusion.

Combining the two components in one reaction required balancing a 1,500 bp target against amplicons of ∼650 bp. Multiplex PCR optimisation demonstrated successful co-amplification of the full-length 16S rRNA target and short AMR amplicons across a range of 16S rRNA primer concentrations, enabling selection of conditions that balanced amplification of long and short targets within a single reaction (**Figure 1d**). The primer concentrations shown in **Figure 1d** represent optimisation experiments used to identify conditions that balanced amplification of long and short targets and do not correspond to the final primer concentrations used in the assay, which employed a non-equimolar primer pool with AMR primer concentrations ranging from 0.4 to 0.8 µM and a 16S rRNA primer concentration of 2.0 µM. Gel profiles showed distinct co-amplification of the short AMR fragments and the long 16S rRNA fragment, with no detectable primer-dimer formation, and nanopore read-length distributions displayed two clear peaks corresponding to the expected AMR and 16S rRNA amplicon sizes, confirming simultaneous amplification (**Figure 1e**). The same reaction tolerated substitution of the ribosomal operon for the 16S gene: all eight AMR amplicons co-amplified with the 4.2–4.5 kb operon in a single tube and again resolved as two distinct read-length populations (**Supplementary Figure S2**). Performance of the complete nine-target assay was then established using the multidrug-resistant clinical isolate pool, which contained panel AMR targets (**Figure 2**).

**Figure 2:**
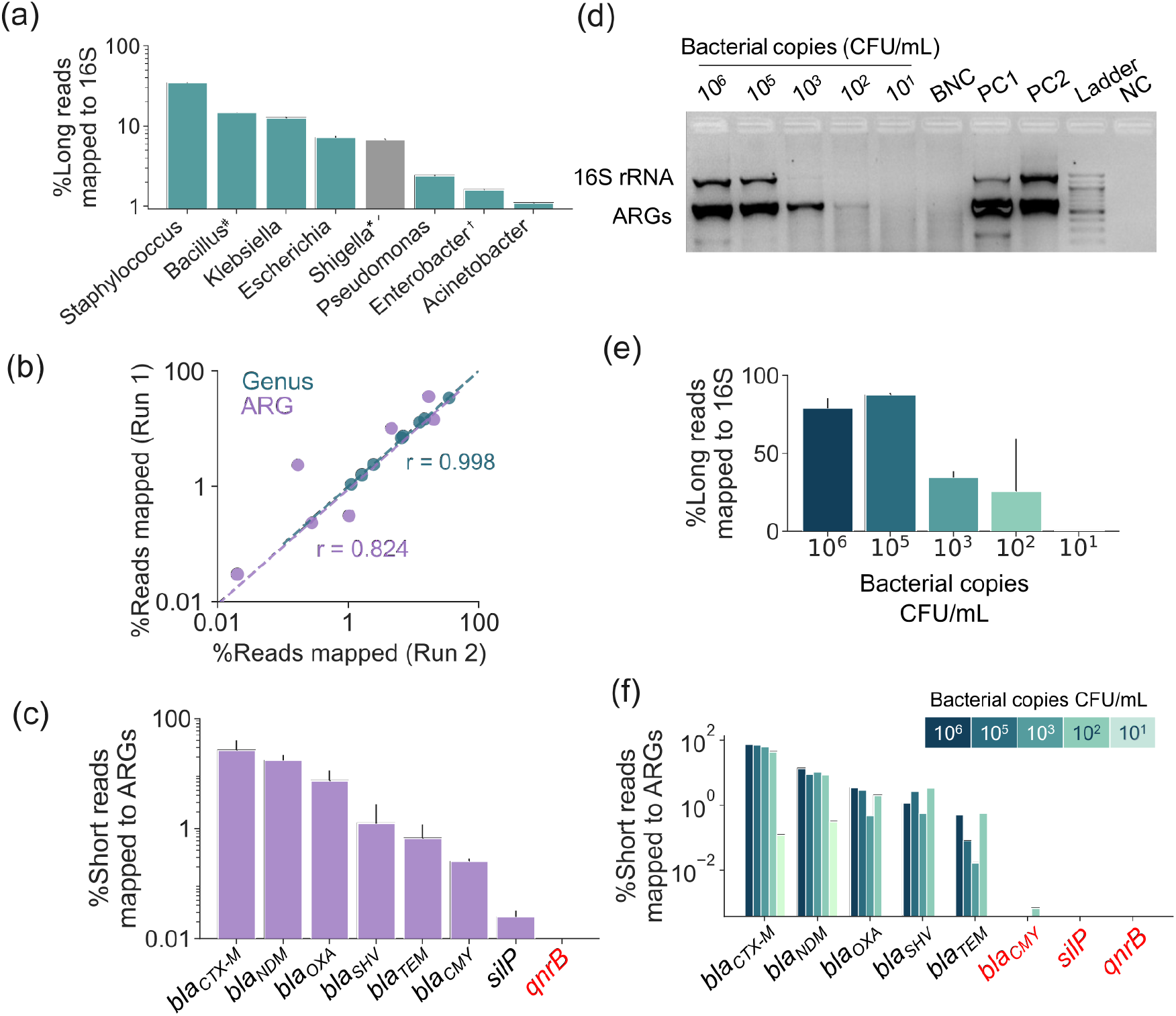
RAMP-Seq performance on complex bacterial samples. **(a)** Genus-level taxonomic assignments of full-length 16S rRNA reads from a pool of multidrug-resistant clinical isolates combined in equal amounts (1 µg genomic DNA per isolate). *Shigella shares >99% 16S rRNA identity with Escherichia. #Bacillus*\* reads derive from *Bacillus safensis* contamination of the isolate DNA preparations, confirmed by Illumina sequencing (**Supplementary Table S8**). †*Enterobacter* was not among the culture-based identifications of the pooled isolates. *Enterococcus*, present in the pool, was detected at 0.06% of classified reads (below the 1% reporting threshold) and is not shown. **(b)** Reproducibility of genus-level (teal) and AMR (purple) profiles between two biological replicates (Pearson’s r = 0.998 for genera and r = 0.824 for AMR targets). **(c)** Panel resistance determinants detected in the same pooled isolates *bla_SHV/LEN/OKP_*, *bla_CMY-6_*, *bla_NDM-1_*, *bla_CTX-M-1_*, *bla_TEM_*, *bla_OXA-48_*, and *silP* were detected; *qnrB1* (red) was not detected by RAMP-Seq, and Illumina WGS identified only the divergent *qnrB4* and *qnrB55* alleles. **(d)** Agarose gel of one-pot multiplex PCR from *Klebsiella pneumoniae* ET/167/2021 spiked into human blood at 10⁶–10¹ CFU/mL. BNC, blood-negative control; PC1, *K. pneumoniae* positive control; PC2, *Escherichia coli* positive control; NC, no-template control. **(e)** Percentage of long reads mapped to the *K. pneumoniae* 16S rRNA gene at each spike concentration. **(f)** Percentage of short reads mapped to each panel AMR target at each spike concentration. Targets in red (*bla*_C*MY-6*_*, silP*, *qnrB1*) were not detected above trace levels by RAMP-Seq (*bla*_C*MY-6*_ was detected in one of the replicates of 10^2^ CFU/mL at 0.0007%) are absent from this isolate by Illumina WGS (**Supplementary Table S8**). Error bars represent the standard deviation of two biological replicates.

### Validation of RAMP-Seq on Multidrug-resistant Clinical Isolates

Having established the taxonomic and resistance components separately, we next asked whether RAMP-Seq resolves all sequences simultaneously in a sample containing both. Genomic DNA (1 µg each) from ten multidrug-resistant clinical isolates spanning six genera—*Escherichia coli* (two isolates), *Klebsiella pneumoniae* (two), *Acinetobacter* spp. (three), *Pseudomonas aeruginosa*, *Enterococcus* spp., and methicillin-resistant *Staphylococcus aureus* (MRSA)—was pooled together with genomic DNA from the *Klebsiella pneumoniae* reference strain MCC 2451 (NCCS/NCMR, Pune) to generate a mixed-community DNA sample, of which 100 ng served as template for the multiplex RAMP-Seq reaction.

Five of the six genera in the pool were detected above the thresholds (>1 %) from full-length 16S rRNA reads **(Figure 2a)**, and taxonomic profiles were highly reproducible across two biological replicates (Pearson’s r = 0.998, **Figure 2b**). Although *Shigella* was not added to the pool, 6.7% of long reads mapped to *Shigella. Escherichia* and *Shigella* are conventionally reported together in nanopore sequencing due to very high similarity (>99%) between the 16S rRNA gene [30].

*Bacillus* accounted for 14.7% of classified reads, a genus absent from the culture-based identifications supplied with the isolates. Illumina sequencing of the individual DNA preparations identified it as *Bacillus safensis.* Reads matched the *B. safensis* 16S rRNA gene with 100% identity (**Supplementary Table S8b**) and covered >90% of the *B. safensis* genome in six of seven preparations (**Supplementary Table S8a**), indicating that the organism itself was present rather than residual DNA. *Bacillus* species are spore-forming and environmentally ubiquitous and are frequently recorded as contaminants of clinical cultures and nucleic-acid reagents [31, 32]. *Enterobacter*, which was not among the culture-based identifications, accounted for ∼1.6% of classified reads; BLAST analysis of a consensus of these reads confirmed *Enterobacter* rather than *Klebsiella*, and its source could not be determined. Conversely, *Enterococcus* (BC/1553/2021-B2), present in the pool and confirmed by Illumina sequencing (**Supplementary Table S8**), was detected at only 0.06% of classified reads, below the reporting threshold.

Seven of the eight ARG panel determinants were detected (**Figure 2c**), and their abundance profiles were reproducible across replicates (Pearson’s r = 0.824, **Figure 2b**). *qnrB1* returned no reads in either biological replicate. Illumina sequencing identified *qnrB4* and *qnrB55* in the pool but not *qnrB1*; these alleles share 86% coding-sequence identity with *qnrB1* and carry four mismatches within the *qnrB1* forward-primer binding site, one of them at the 3′ position, so they are not expected to prime. Detection was therefore complete in both directions; every panel target present in the pool was detected, and the single panel target absent from it generated no reads, indicating that the *qnrB1* primers do not cross-amplify these divergent alleles and the *qnrB1* channel served as an internal control for allele-level specificity, confirming that the primers do not cross-amplify divergent members of the same gene family. Whole-genome sequencing was performed on seven of the ten isolates (**Supplementary Table S8**); the remaining three were not sequenced, so concordance is assessed against the determinants identified in those seven.

Together, the taxonomic and resistance profiles show that RAMP-Seq resolves both the composition of a complex clinical mixture and its panel resistance content from a single reaction, reproducibly across independent runs.

### Analytical Sensitivity in Spiked Human Blood

To assess analytical sensitivity in a clinically relevant matrix, *Klebsiella pneumoniae* ET/167/2021-B1 was spiked into human blood at 10⁶–10¹ CFU/mL and carried through the complete workflow, including carbonate-based host DNA depletion, whose performance was first established with Escherichia coli-spiked blood **(Supplementary Figure S3)**. Multiplex amplicons were recovered across the titration series **(Figure 2d).**

Under the conditions tested, 10² CFU/mL represented the lowest concentration at which *Klebsiella pneumoniae* was detected in both biological replicates, with both culture and sequence confirmation (**Figure 2e**). Detection of *Klebsiella pneumoniae* was not reproducibly observed at the next lower concentration tested (10 CFU/mL). Because only two biological replicates were analysed at each dilution, this value is reported as the lowest reproducibly detected concentration rather than a formally validated limit of detection. Replicate-level sequencing statistics are provided in **Supplementary Tables S7 and S9**. The fraction of unmapped reads correlated strongly and inversely with bacterial load (Pearson’s r = −0.939; **Supplementary Figure S5**), suggesting that this metric may serve as a qualitative biomarker of low biomass.

Panel resistance determinants were recovered across the titration series (**Figure 2f**). *bla_CTX-M_* and *bla_NDM_* were detected at every concentration tested, down to 10¹ CFU/mL, while *bla_OXA_*, *bla_SHV/LEN/OKP_* and *bla_TEM_* were detected from 10⁶–10^2^ CFU/mL. *qnrB1* and *silP* were not detected above trace levels at any concentration while *bla*_C*MY-6*_ was detected in one of the replicates of 10^2^ CFU/mL at 0.0007%, consistent with their absence from this isolate by Illumina whole-genome sequencing (**Supplementary Table S8**). Resistance determinants were therefore detectable at or below the load at which the organism itself was reproducibly identified, which is the behaviour required of a workflow intended to report both from one reaction.

Separate exploratory experiments tested whether the longer ribosomal target performs in the same matrix. *Escherichia coli* spiked into human blood yielded long reads mapping to the expected 16S-ITS-23S rRNA operon (∼4.2–4.5 kb) at 10⁵, 10³ and 10¹ CFU/mL **(Supplementary Figure S6).** These experiments evaluated compatibility with longer ribosomal targets and were not used to determine analytical sensitivity, detection limits, or overall workflow performance.

### Application of RAMP-Seq to Wastewater and Surface Water

The mock community, isolate pool and blood spikes each had a defined expected composition. We next applied the complete nine-target reaction to water samples, where the community is unknown and the matrix varies. Samples were collected from five sites spanning a wastewater treatment-to-freshwater gradient in Bengaluru, India (**Figure 3a; Supplementary Table S2**). This comprised of influent (RC_In_) and effluent (RC_Out_) from the Raja Canal sewage treatment plant (STP), and since treated effluent from this and neighbouring STPs discharges into Bagalur Lake, the lake inlet (BL_In_), the lake outlet (BL_Out_), and a nearby open well (BL_OW_). These matrices vary in turbidity, microbial biomass, and community complexity and collectively assess whether the workflow can produce interpretable taxonomic and resistance profiles beyond predefined mixtures.

**Figure 3:**
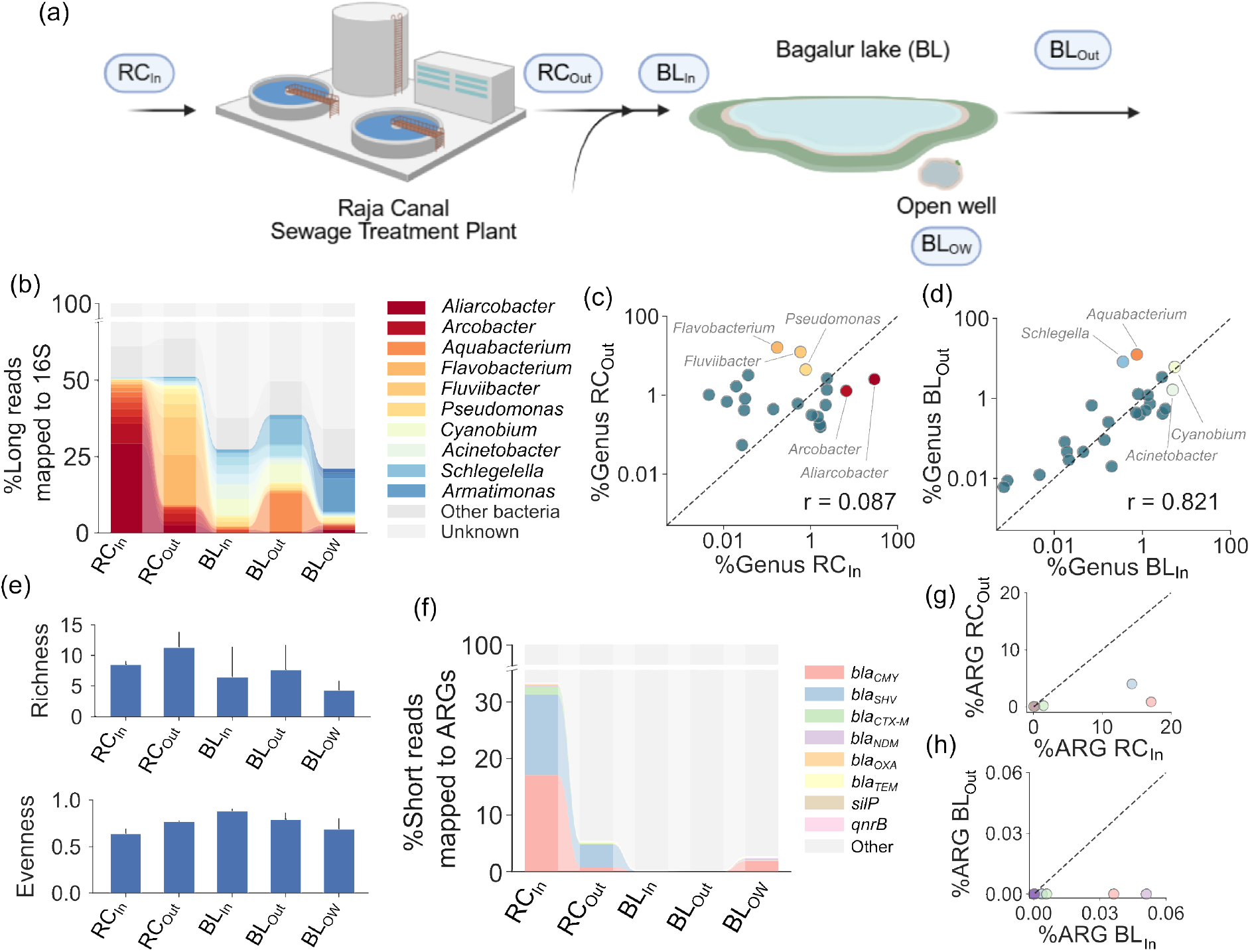
Environmental and wastewater surveillance using RAMP-Seq. **(a)** Schematic depicting sample collection sites across the Raja Canal sewage treatment plant and Bagalur Lake system. **(b)** Relative abundances of bacterial genera across all sites; the legend shows the top 10 genera detected across all samples. Genera with <1% relative abundance were grouped as “other bacteria”. **(c)** Correlation of genus-level taxonomic profiles between Raja Canal STP influent (RC_In_) and effluent (RC_Out_). (Pearson’s r = 0.087). **(d)** Correlation of genus-level taxonomic profiles between Bagalur Lake inlet (BL_In_) and outlet (BL_Out_) (Pearson’s r = 0.821). **(e)** Richness and evenness of samples collected at each site. **(f)** Distribution of targeted AMR determinants across all sampling sites. **(g)** Correlation of targeted AMR profiles between Raja Canal STP influent (RC_In_) and effluent (RC_Out_). **(h)** Correlation of targeted AMR profiles between Bagalur Lake inlet (BL_In_) and outlet (BL_Out_). RC_In_, Raja Canal STP influent; RC_Out_, Raja Canal STP effluent; BL_In_, Bagalur Lake inlet; BL_Out_, Bagalur Lake outlet; BL_OW_, Bagalur Lake open w

Influent wastewater samples were predominantly composed of members of the family Arcobacteraceae, which constituted 35.8% of classified reads. These taxa were detected under both current and former genus-level annotations (*Aliarcobacter*, 29.2%; *Arcobacter*, 6.6%) (**Figure 3b**), consistent with their known prevalence as enteric organisms in untreated sewage [33]. In contrast, effluent samples were enriched for *Flavobacterium* (16.1%), an aerobic degrader of complex organic matter abundant in activated sludge, and *Fluviibacter* (12.4%), a polyphosphate-accumulating freshwater bacterioplankton (**Figure 3b–c**) [34, 35]. Genus-level profiles of influent and effluent were essentially uncorrelated (Pearson’s *r* = 0.087; **Figure 3c**). Detailed genus-level profiles for all environmental samples are provided in **Supplementary Table S10**.

The Bagalur Lake inlet yielded fewer classified reads than the STP effluent and displayed a shift towards photoautotrophic genera such as *Cyanobium* and *Synechococcus*. The lake outlet showed a similar community composition to the inlet (Pearson’s r = 0.821; **Figure 3d**) and contained *Aquabacterium*, a taxon commonly associated with freshwater biofilms. The open well exhibited the most distinct microbial profile, with approximately 10% of reads assigned to *Armatimonas*. Although overall richness was similar across samples (**Figure 3e**), the Bagalur Lake inlet exhibited the most even taxonomic distribution, whereas the STP samples were dominated by a smaller number of highly abundant taxa. Genus-level profiles were reproducible between biological replicates at every site (**Supplementary Figure S7**).

AMR profiling revealed multiple targeted resistance determinants in influent wastewater, with *bla_CMY-6_* and *bla_SHV/LEN/OKP_* being the most abundant panel targets, both associated with β-lactam resistance (**Figure 3f**). Effluent samples exhibited lower relative abundances of panel determinants than influent (**Figure 3g**), and the lake and well samples contained few or no detectable panel targets (**Figure 3f, h**). Accordingly, the panel determinants were detected against a background comprising several hundred genera with relative abundances well below 1%, while no detectable signal was observed in the least impacted matrices. These results indicate that the panel provides the requisite dynamic range and analytical specificity for application to complex environmental mixtures.

Across all five matrices, the workflow produced classifiable full-length 16S profiles and panel resistance readouts from the same reaction, separating samples that differ markedly in biomass, turbidity and community structure. Because RAMP-Seq quantifies the relative abundances of selected targets, inter-site variation is presented here in terms of differences in targeted read profiles, and their interpretation in an environmental context is addressed in the Discussion.

## Discussion

RAMP-Seq was developed to address a long-standing challenge in AMR surveillance, namely the need to combine taxonomic identification with resistance gene detection in a single, rapid, and operationally simple workflow. Long-read nanopore sequencing has already found an important niche in clinical and environmental AMR surveillance owing to its portability, real-time data generation, and flexibility in assay development [36], and recent nanopore-based assays have demonstrated that multiplex PCR coupled with nanopore sequencing can enable same-day identification of pathogens and resistance determinants from bloodstream infections [13, 37]. Such targeted workflows typically provide greater sensitivity for pathogen and AMR detection than unbiased metagenomic sequencing, particularly in low-biomass clinical samples; however, they remain constrained by reliance on predefined target panels and often require separate amplification workflows for taxonomic and resistance profiling [38, 39]. RAMP-Seq builds on this foundation by integrating long- and short-amplicon targets within a single reaction so that one amplification and one barcode per sample yield both profiles, distinguishing it from approaches that rely on multiple primer pools or separate reactions and library preparations [13].

Full-length 16S rRNA profiling gave reproducible genus-level profiles across the mock community and the clinical isolate pool. As with any 16S rRNA-based method, closely related taxa such as *Escherichia* and *Shigella* cannot be reliably separated [30], so we report them conservatively at the *Escherichia*/*Shigella* group level. However, two discrepancies with the culture-based composition of the clinical pool were observed: *Enterococcus*, although present, was recovered at only 0.06% of classified reads, below the reporting threshold, and *Enterobacter* reads were detected although no *Enterobacter* isolate was included. The low Enterococcus recovery may result from multiplex amplification bias against this template, suggesting genus-level recovery from mixed templates can be incomplete.

Resistance detection was concordant with Illumina whole-genome sequencing of seven clinical isolates (**Supplementary Table S8**). Every panel determinant present in the pool was detected, and the one panel target absent from it, *qnrB1*, returned no reads. The isolates carried *qnrB4* and *qnrB55*, which differ from *qnrB1* at the forward-primer binding site and were not expected to amplify. Thus, the qnrB primers distinguish alleles but restrict detection to alleles matching the primer site: isolates with divergent qnrB alleles will be missed because RAMP-Seq discriminates at primer-binding sites, not within-amplicon variants.

A central technical challenge addressed by RAMP-Seq is the need to perform multiplex PCR across a wide amplicon size range. Simultaneous amplification of multiple targets leads to primer competition and reduced amplification efficiency compared with singleplex PCR assays, and co-amplification of the ∼1.5 kb 16S rRNA gene alongside multiple short AMR targets introduces competition for reaction components and an amplification bias that favours the shorter fragments [40, 41]. Rather than optimising the sensitivity of any single target, RAMP-Seq was designed to achieve balanced and reproducible recovery of both long and short amplicons within a single tube, accepting reduced analytical sensitivity relative to singleplex assays in exchange for analytical integration and interpretability. Advances in nanopore chemistry and basecalling have further strengthened such workflows: evaluations of the R10.4.1 chemistry have demonstrated improved consensus accuracy and reliable genome reconstruction from long-read data [42]. Consistent with other nanopore-based targeted sequencing approaches, relatively permissive alignment thresholds were employed to accommodate residual sequencing errors while maintaining detection performance.

Analytical sensitivity experiments using spiked human blood reflect these constraints. RAMP-Seq reproducibly detected bacterial targets down to 10² CFU/mL following host DNA depletion under the experimental conditions evaluated. Although bloodstream infections may occur at concentrations below 10 CFU/mL [43], loads of 10²–10³ CFU/mL are frequently reported in paediatric sepsis, early-stage infection and post-enrichment diagnostics [43–45], and the value obtained here is comparable to the detection limit of approximately 90 CFU/mL reported for nanopore full-length 16S rRNA gene sequencing of normally sterile body fluids [39]. Clinical detection performance is nevertheless likely to vary according to pathogen burden, host DNA background, sample quality, and specimen-processing workflows.

Application of RAMP-Seq to wastewater and downstream freshwater samples highlights its relevance within a One Health surveillance framework. Between sewage treatment plant influent and effluent, the genus-level community was almost entirely replaced (Pearson’s r = 0.087), and both the number and the relative abundance of panel determinants detected fell markedly, with few or no panel targets recovered from the downstream lake and well samples. Comparable shifts in community composition and resistance-gene content across wastewater treatment have been reported in metagenomic and nanopore-based resistome studies of sewage influent and effluent [46–48], and long-read resistome profiling has further demonstrated the persistence and mobility of resistance determinants across wastewater systems [49]. RAMP-Seq captures these patterns using a targeted workflow that requires substantially lower sequencing depth and computational resources than shotgun metagenomics.

The eight-target panel was selected for public health importance, particularly in the Indian AMR landscape, where ESBLs, carbapenemases, quinolone-resistance determinants, and AmpC β-lactamases are prevalent across clinical and environmental samples and are frequently associated with mobile genetic elements [15–18]. Future iterations may incorporate additional clinically important resistance determinants, including further carbapenemases, colistin resistance, and Gram-positive resistance markers, as multiplex optimisation strategies continue to improve. Pilot amplification of the full 16S–ITS–23S rRNA operon further demonstrates compatibility with longer ribosomal targets and suggests a pathway to improved phylogenetic resolution in future iterations, as shown in recent long-read rRNA sequencing studies [50]. The complete workflow, from extracted DNA through multiplex PCR, nanopore library preparation, sequencing, and preliminary analysis, was completed in 8.2 ± 0.6 hours; this same-day turnaround may be particularly valuable in settings with a high burden of AMR and limited laboratory infrastructure.

Several considerations apply when interpreting these findings. The study was designed as a technical validation of a unified taxonomic and resistance profiling workflow and used a limited number of biological replicates across multiple sample types. Analytical performance in blood was evaluated using contrived human blood specimens spiked with cultured bacterial isolates rather than prospectively collected clinical samples. Environmental AMR abundance was inferred from targeted amplicon sequencing rather than direct quantification of absolute gene copy number, so differences between sampling locations reflect relative abundances within individual libraries. Because environmental samples were collected during a single sampling event, replicate measurements represent independent DNA extractions and amplifications from the same grab sample rather than independent sampling events. Detection of resistance genes reflects resistance potential and may not always correspond to expressed phenotypic resistance. Future studies involving larger sample sets, prospective clinical specimens, expanded target panels, repeated environmental sampling, and longitudinal monitoring, combined with quantitative approaches such as qPCR or digital PCR, will further define the applicability of RAMP-Seq across One Health surveillance settings. RAMP-Seq is not intended to replace comprehensive resistome or genome-wide analyses but to provide a rapid, integrated snapshot of microbial identity and high-priority resistance determinants, positioning it as a practical complement to deep metagenomic sequencing within a One Health framework.

## Conclusion

Across a defined mock community, a pooled set of multidrug-resistant clinical isolates, bacteria spiked into human blood, and wastewater and surface-water matrices, RAMP-Seq yielded full-length 16S rRNA–based taxonomic profiles and targeted antimicrobial-resistance gene readouts from the same single-tube amplification. Detection was reproducible across independent replicates and, for all panel determinants detected, demonstrated concordance with whole-genome sequencing. Prospective evaluation in clinically collected specimens, together with expansion of the panel to include additional alleles within the targeted gene families, constitutes the logical next phase of validation and assay development.

## Author Contributions

- **Deepali Pathak** conceived and designed the study, performed multiplex PCR and Nanopore sequencing, and contributed to manuscript writing.
- **Suraj Jagtap** analysed sequencing data, prepared figures, assisted with data interpretation, and contributed to manuscript writing.
- **Rekha Godbole** isolated genomic DNA and plasmids from AMR and wastewater bacterial samples and assisted with human-spiked blood DNA experiments.
- **Nirali Patel** designed and coordinated environmental sampling, optimized wastewater sample processing and DNA extraction workflows, and performed wastewater sample pre-processing.
- **Ameena Mohammed** contributed to antimicrobial resistance (AMR) primer design and PCR validation.
- **Jeevanandan Sree Priyanka** assisted with primer optimisation, Sanger sequencing, and Nanopore standardisation.
- **Debalina Sarkar** conducted spike-in blood experiments and wastewater PCR assays.
- **Imon Roy** and **Varsha Sridhar** contributed to environmental sample collection and DNA preparation. Varsha Sridhar also supported coordination of the wastewater surveillance studies.
- **Arvind Natarajan** provided clinical isolates essential to the study.
- **Vishwanath Srikantaiah** provided domain expertise and strategic guidance on water systems and the study’s sustainability aspects.
- **Rahul Roy** conceived, designed and supervised the study and contributed to experimental design, planning, and manuscript writing.

## Acknowledgements

The authors thank Saranya Marimuthu for her assistance and valuable feedback on this work. The content is solely the responsibility of the authors and does not necessarily represent the official views of the National Institutes of Health.

## Abbreviations

AMR: Antimicrobial resistance
PCR: Polymerase chain reaction
ONT: Oxford Nanopore Technologies
WGS: Whole-genome sequencing
RAMP-Seq: Rapid AMR Profiling by Sequencing

## Supplementary Figures

**Supplementary Figure S1:**
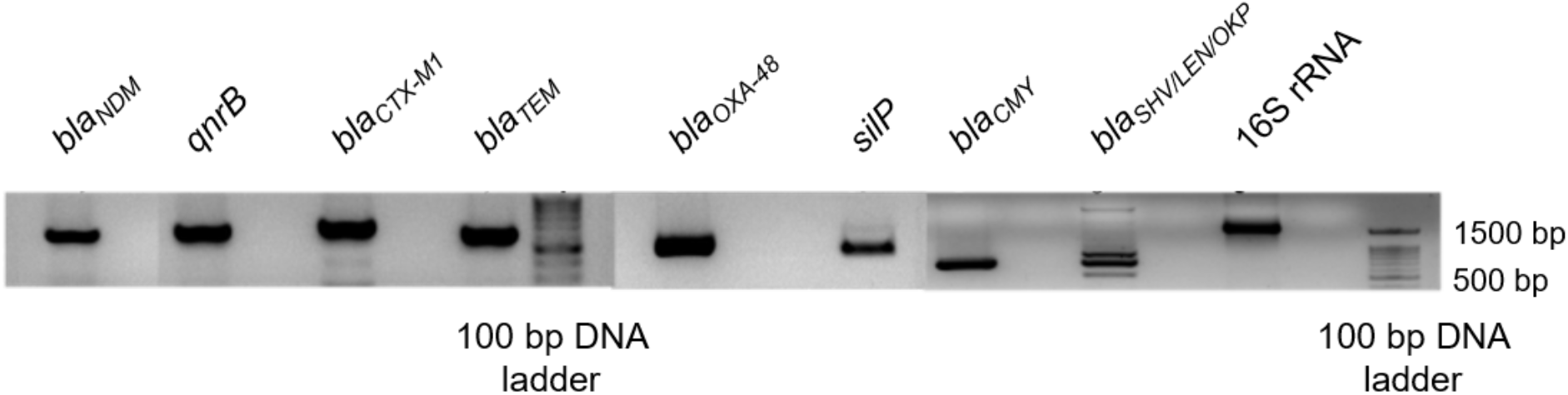
Agarose gel electrophoresis showing PCR amplification of antimicrobial resistance (AMR) genes and the 16S rRNA gene. The 100 bp DNA ladder is shown for size reference; amplified products range from ∼600 bp to ∼1,500 bp, corresponding to the expected sizes of the target genes.

**Supplementary Figure S2:**
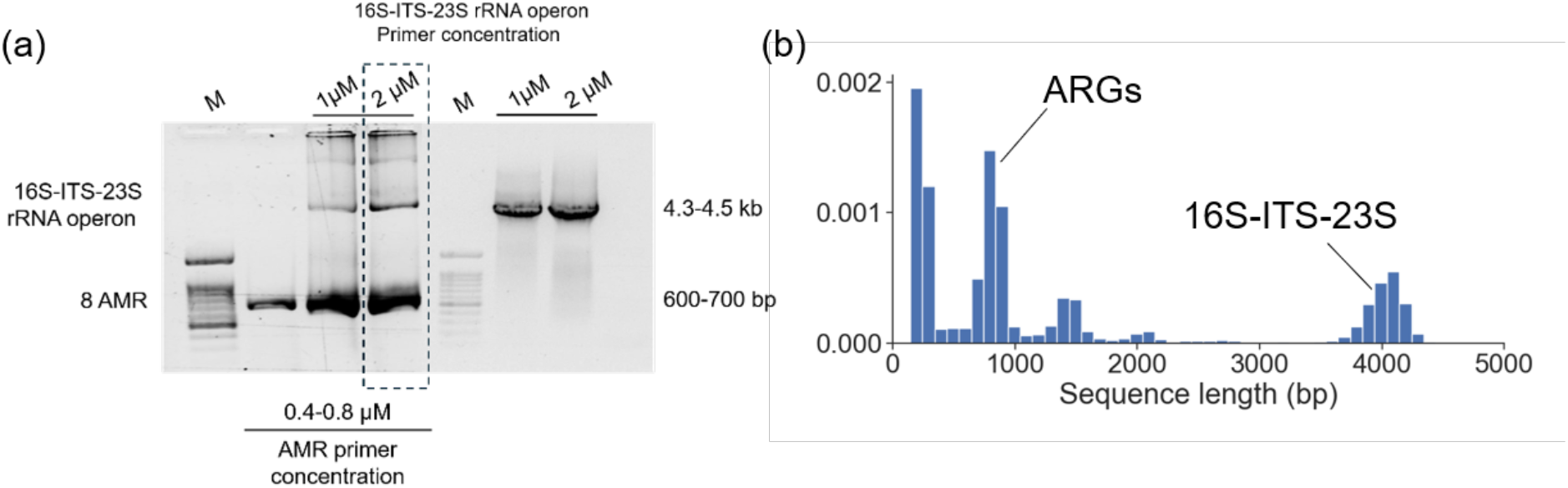
Co-amplification of ARGs and 16S-ITS-23S gene amplicons **(a)** Evaluation of long-range ribosomal operon and AMR gene co-amplification in a multiplex PCR assay. Multiplex PCR reactions were performed to assess the compatibility of simultaneous amplification of eight AMR gene targets and the long-range 16S-ITS-23S rRNA operon. Reactions containing optimised AMR primer concentrations (0.4-0.8 µM) were tested with 16S-ITS-23S rRNA primers at 1 µM and 2 µM. Agarose gel electrophoresis revealed successful amplification of the ∼4.2–4.5 kb ribosomal operon alongside AMR gene amplicons ranging from ∼600 to 700 bp. The assay demonstrated efficient co-amplification of both long and short targets within a single reaction, with the optimised primer concentrations providing balanced amplification and supporting their use in the final multiplex PCR workflow. **(b)** The read length distribution shows dominant peaks corresponding to ARGs and 16S-ITS-23S rRNA amplicons.

**Supplementary Figure S3:**
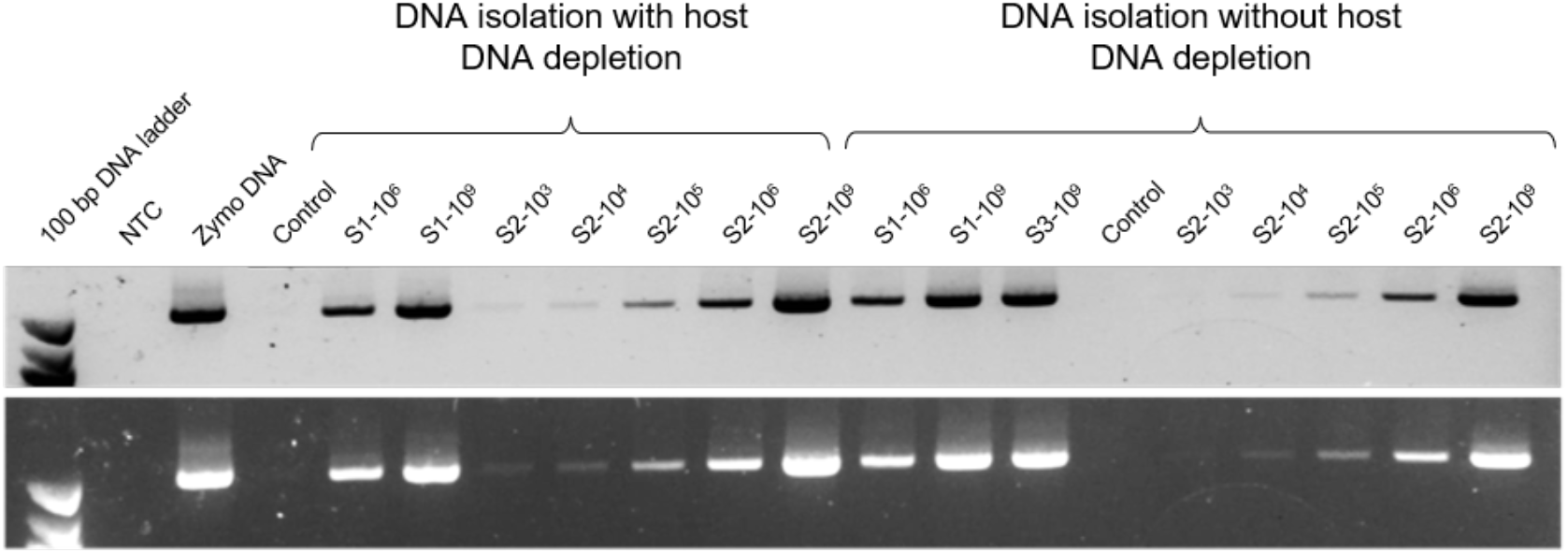
Effect of carbonate-based host DNA depletion on 16S rRNA amplification from *Escherichia coli*-spiked human blood. Three independent human blood samples (S1–S3; 1 mL each) were spiked with *Escherichia coli* at 10³– 10⁹ CFU/mL, and DNA was extracted either with carbonate-lysis host DNA depletion or directly without depletion. Lane labels give the blood sample and spike concentration (e.g., S2-10⁵ = blood sample 2 spiked at 10⁵ CFU/mL). The full-length 16S rRNA gene (∼1.5 kb) was amplified with 27F/1492R primers. Lowercase “control”, unspiked blood processed with the corresponding extraction method; “NTC”, no-template control; Zymo DNA, ZymoBIOMICS DNA standard (positive control); 100 bp, DNA ladder. Upper and lower panels show the same gel at two exposures.

**Supplementary Figure S4:**
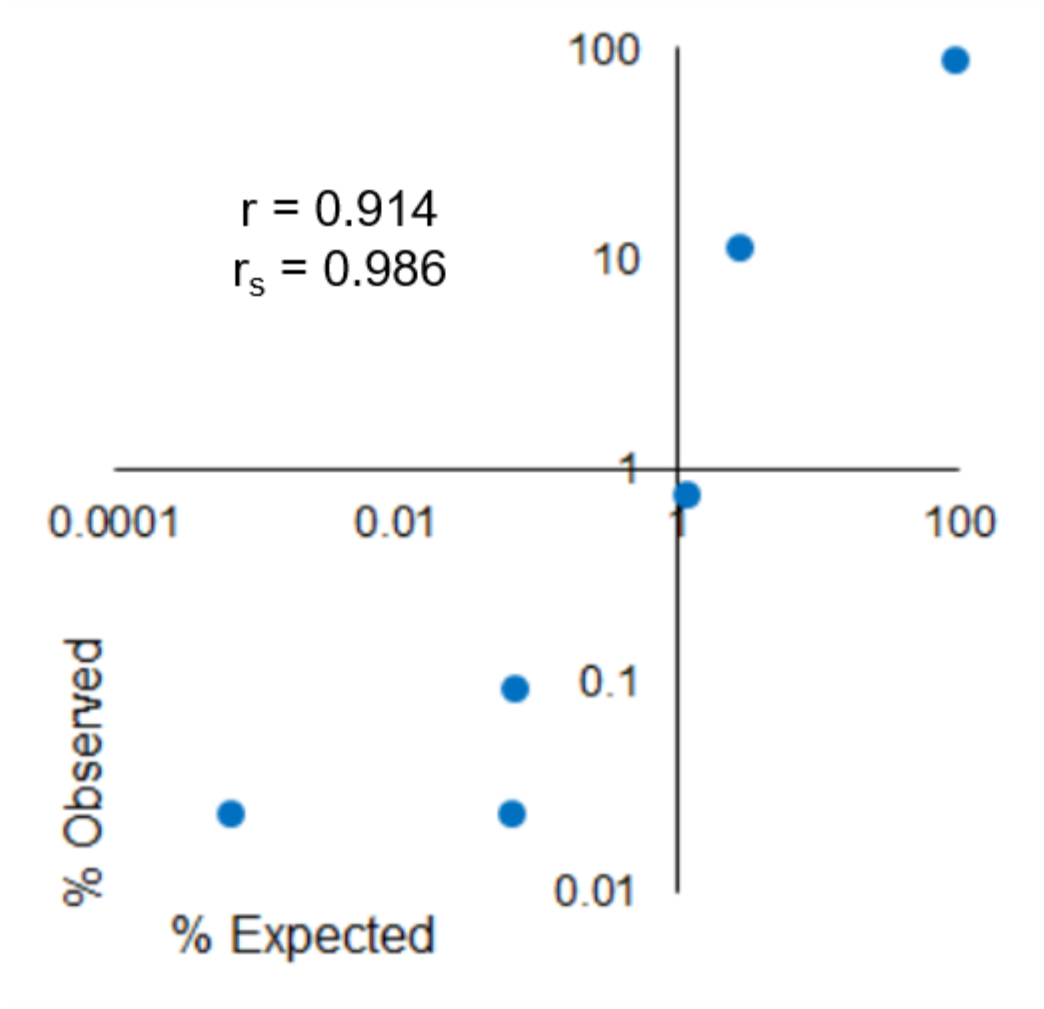
Relative abundance of full-length sequences of 16S-ITS-23S rRNA operon from an 8-member mock microbial community with a log abundance distribution. (Pearson’s r = 0.914, Spearman’s r_s_ = 0.986)

**Supplementary Figure S5:**
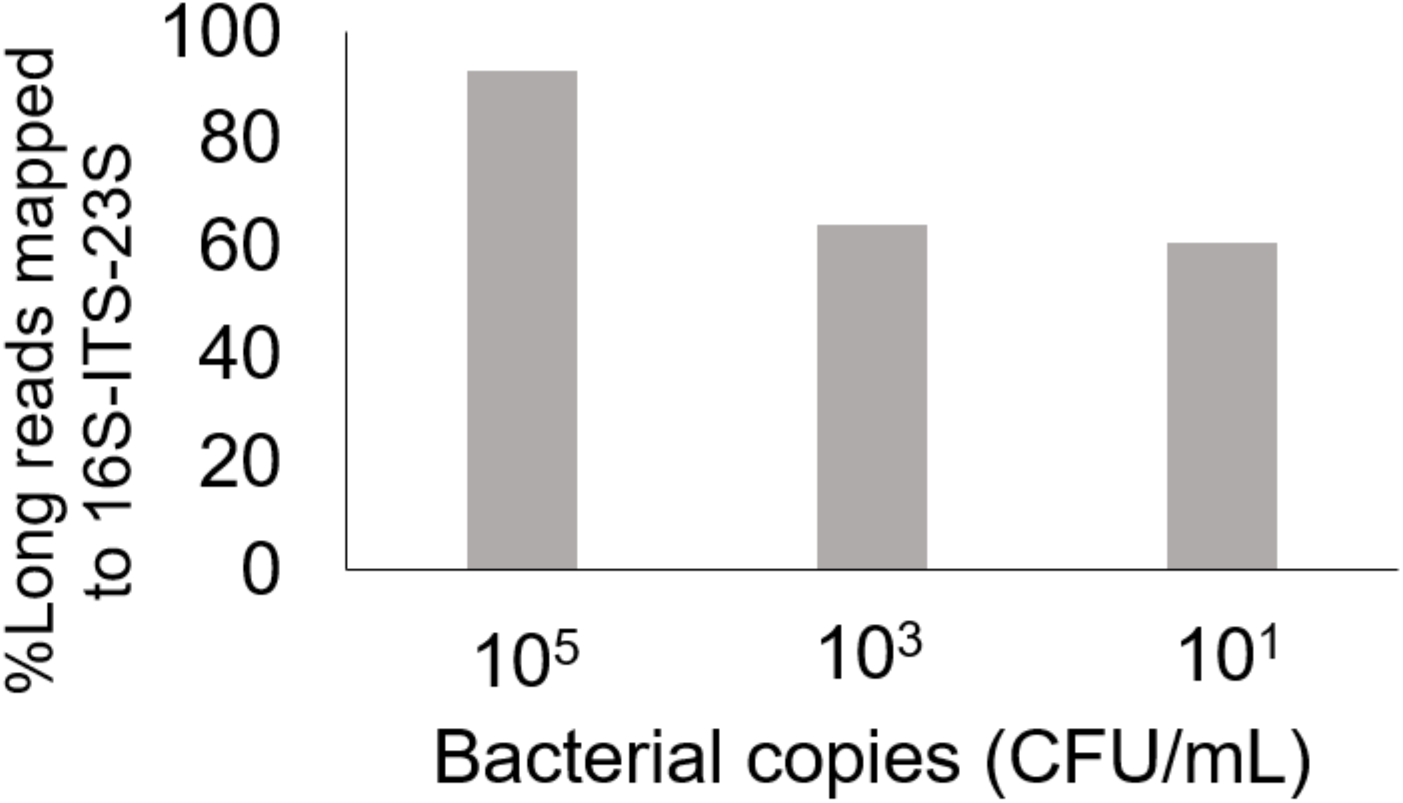
The percentage of unmapped reads shows a negative correlation with bacterial load. Bars show the unmapped read fraction across 10¹–10⁵ CFU/mL (mean ± SD). The unmapped read percentage decreases as CFU increases, establishing unmapped reads as a qualitative indicator of low microbial biomass. (Pearson’s r = −0.939)

**Supplementary Figure S6:**
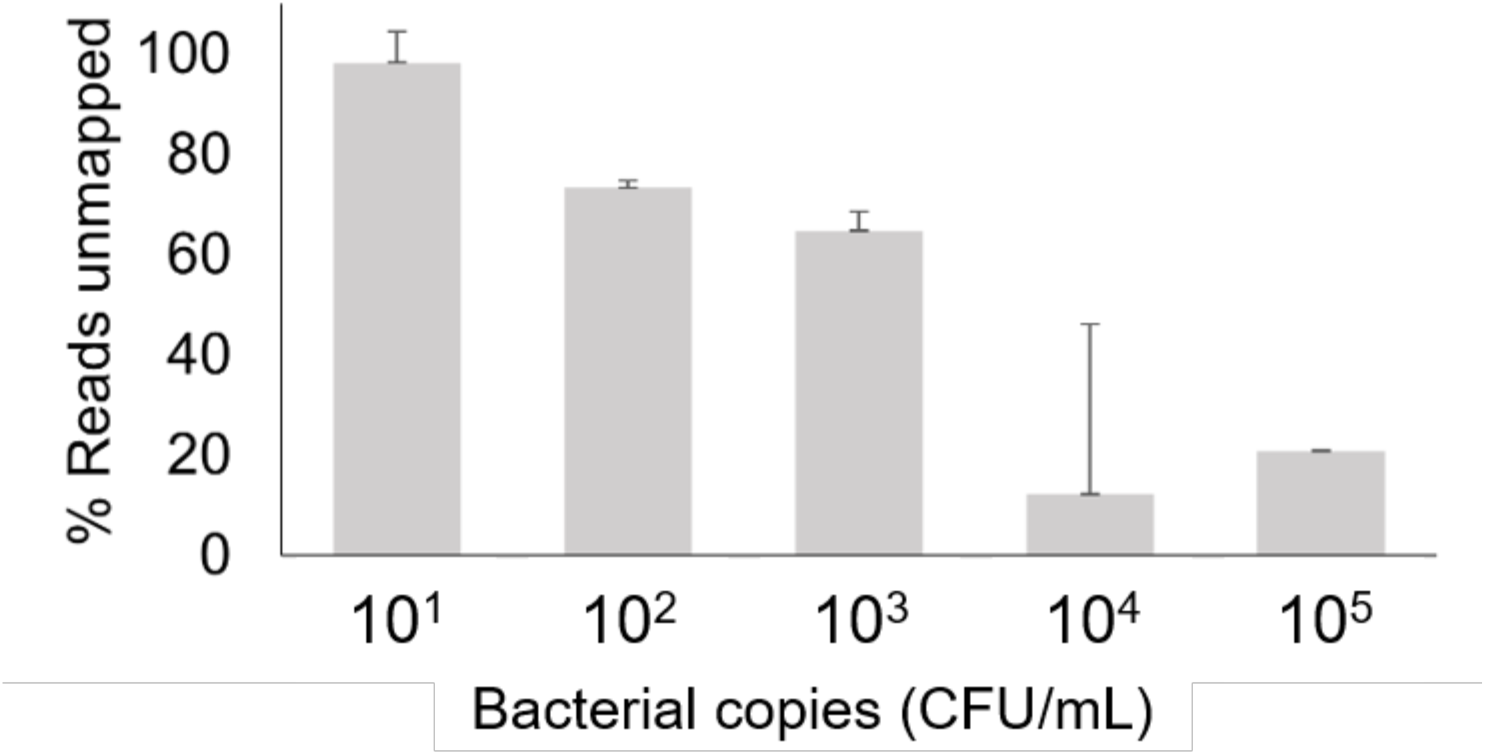
Exploratory evaluation of 16S-ITS-23S operon amplification using *Escherichia coli* spiked into human blood. Percentage of long reads (3000-5000 bp) mapping to the expected 16S-ITS-23S operon amplicon at 10⁵, 10³, and 10¹ CFU/mL. These experiments were performed to assess compatibility of the RAMP-Seq workflow with longer ribosomal targets and were not included in the formal analytical sensitivity evaluation.

**Supplementary Figure S7:**
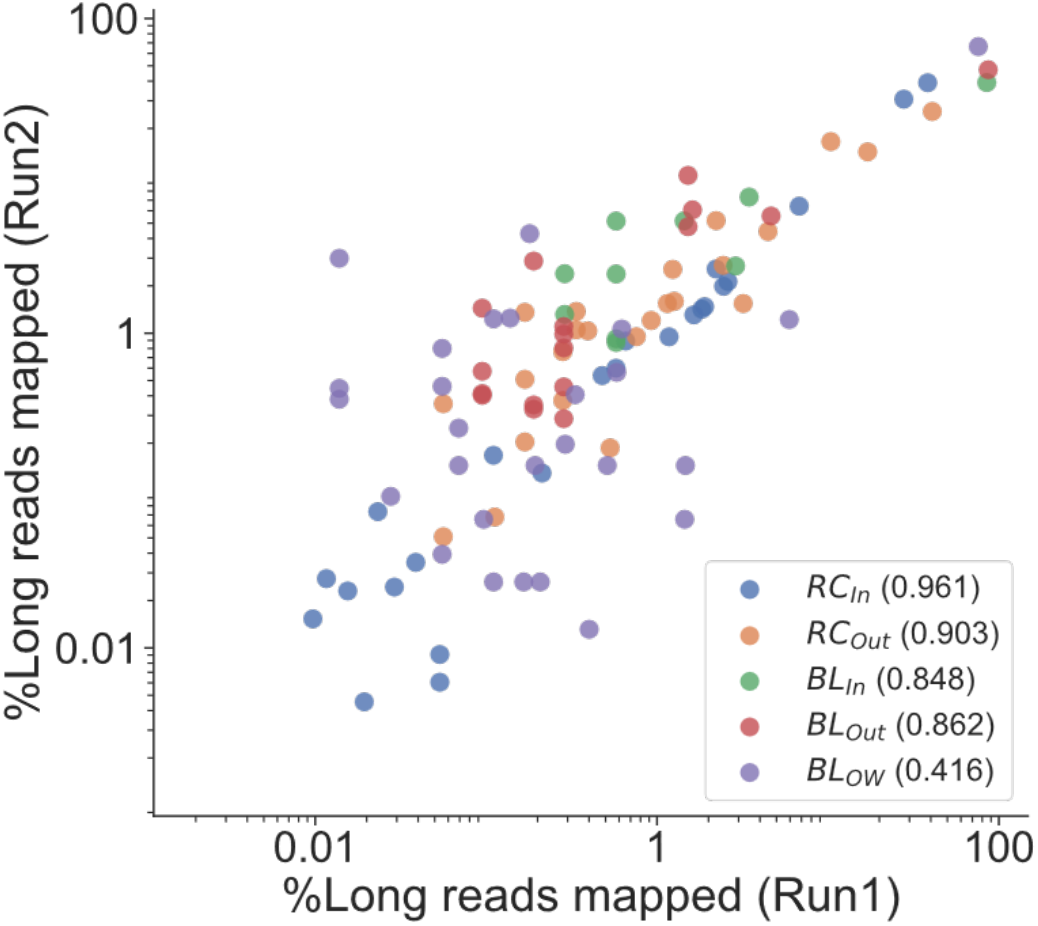
Reproducibility in identifying the genus between the biological replicates of environmental samples. The Pearson’s r is reported for each collection site.

**Supplementary Table S1.** Antimicrobial susceptibility profiles of clinical isolates (Kirby–Bauer disc diffusion) Source: CDLS Microbiology culture reports (2021).

| SN | Lab ID | Sample | Organism | Sensitive (S) | Resistant (R) |
|---|---|---|---|---|---|
| 1 | BC/1553/2021 | Blood culture | *Escherichia coli* BC/1553/2021-B1 | None | AMP, AMC, PIT, CTX, CTR, CAZ, IPM, MRP, ETP, GEN, TOB, AK, CIP, LE, DO, C, COT |
| 2 |  |  | *Enterococcus* spp. BC/1553/2021-B2 | C, VA, LZ | AMP, P, HLG, LE, TE, E |
| 3 | ET/167/2021 | Endotracheal aspirate | *Klebsiella pneumoniae* ET/167/2021-B1 | None | AMP, AMC, PIT, CTX, CTR, CAZ, IPM, MRP, ETP, GEN, TOB, AK, CIP, LE, DO, C, COT |
| 4 | ET/168/2021 | Endotracheal aspirate | *Acinetobacter* spp. ET/168/2021-B1 | None | A/S, PIT, CTR, CAZ, CPM, IPM, MRP, GEN, TOB, AK, CIP, LE, DO, C, COT |
| 5 | P/617/2021 | Pus | *Escherichia coli* P/617/2021-B1 | GEN, TOB, AK | AMP, AMC, PIT, CTX, CTR, CAZ, IPM, MRP, ETP, CIP, LE, DO, C, COT |
| 6 |  |  | *Pseudomonas aeruginosa* P/617/2021-B2 | None | PI, PIT, CAZ, CPM, IPM, MRP, GEN, TOB, AK, CIP, LE |
| 7 | ET/164/2021 | Endotracheal aspirate | *Acinetobacter* spp. ET/164/2021-B1 | A/S | PI, PIT, CTR, CAZ, CPM, IPM, MRP, GEN, TOB, AK, CIP, LE, DO, COT |
| 8 | SP/174/2021 | Sputum | *Klebsiella pneumoniae* SP/174/2021-B1 | TE | AMP, AMC, PIT, CTX, CTR, CAZ, IPM, MRP, ETP, GEN, TOB, AK, CIP, LE, DO, C, COT |
| 9 | NS/315/2021 | Nasal swab | *Staphylococcus aureus* (MRSA) NS/315/2021-B1 | C, COT, LZ | P, AMC, GEN, CIP, TE, E, CD |
| 10 | BC/1559/2021 | Blood culture | *Acinetobacter* spp. BC/1559/2021-B1 | None; Colistin pending (E-strip) | PI, A/S, PIT, CTR, CAZ, CPM, IPM, MRP, GEN, TOB, AK, CIP, LE, DO, COT |

**Abbreviations:** AMP, ampicillin; P, penicillin; PI, piperacillin; AMC, amoxicillin-clavulanate; A/S, ampicillin-sulbactam; PIT, piperacillin-tazobactam; CTX, cefotaxime; CTR, ceftriaxone; CAZ, ceftazidime; CPM, cefepime; IPM, imipenem; MRP, meropenem; ETP, ertapenem; GEN, gentamicin; HLG, high-level gentamicin; TOB, tobramycin; AK, amikacin; CIP, ciprofloxacin; LE, levofloxacin; DO, doxycycline; TE, tetracycline; E, erythromycin; CD, clindamycin; C, chloramphenicol; COT, co-trimoxazole; VA, vancomycin; LZ, linezolid; MRSA, *methicillin-resistant Staphylococcus aureus*; BLI, β-lactamase inhibitor.

**Supplementary Table S2:** Environmental sampling sites around Raja Canal sewage treatment plant and Bagalur Lake.

| **SN** | **Site** | **GPS coordinates** | **Date of sample collection** | **Sample type** |
|---|---|---|---|---|
| 1 | Raja Canal STP | 13.043696971137004, 77.64624863077543 | August 30, 2023 | Influent |
| 2 | Raja Canal STP | 13.043696971137004, 77.64624863077543 | August 30, 2023 | Effluent |
| 3 | Bagalur Lake | 13.138331670665602, 77.66438600674246 | August 30, 2023 | Inlet |
| 4 | Bagalur Lake | 13.141988443107953, 77.66554472096828 | August 30, 2023 | Outlet |
| 5 | Bagalur open well | 13.138051547917607, 77.65731509910577 | August 30, 2023 | Well |

**Supplementary Table S3:**
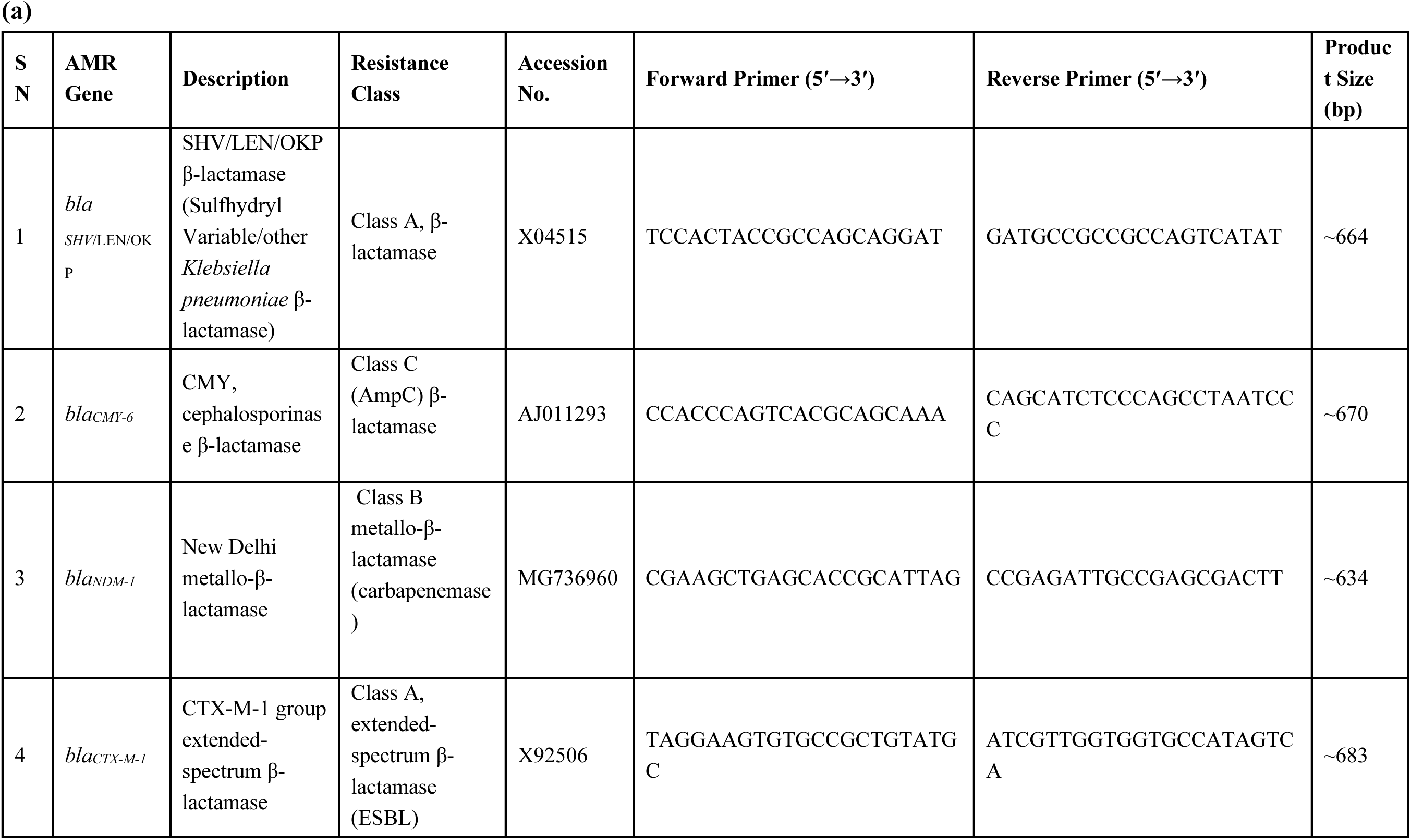
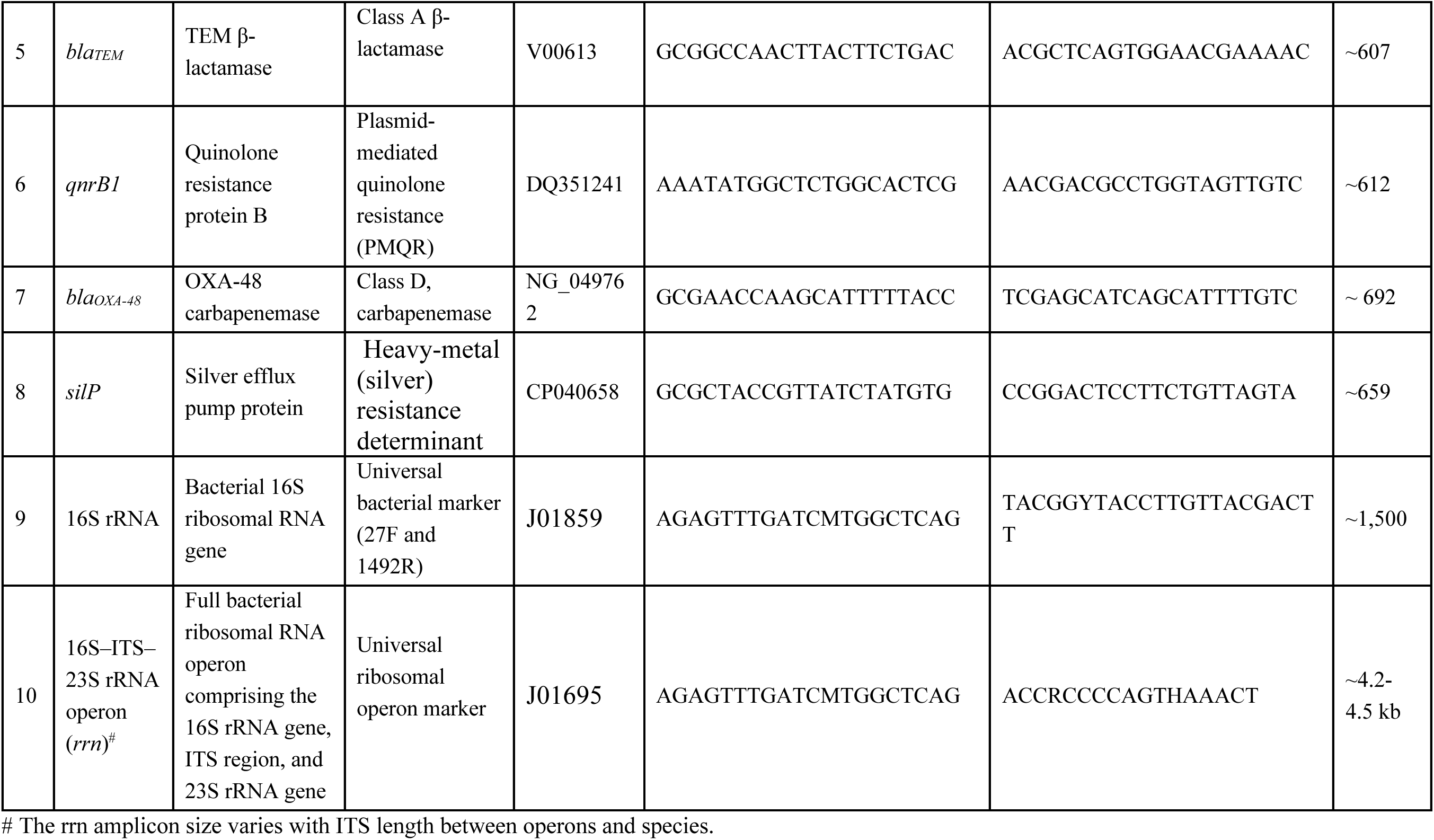
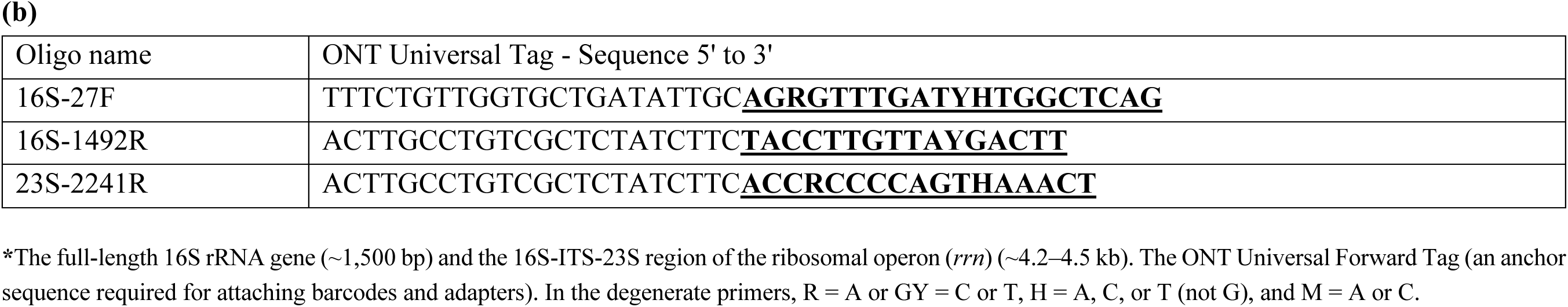
Primer sequences, target genes, and expected amplicon sizes used in RAMP-Seq.

**Supplementary Table S4:**
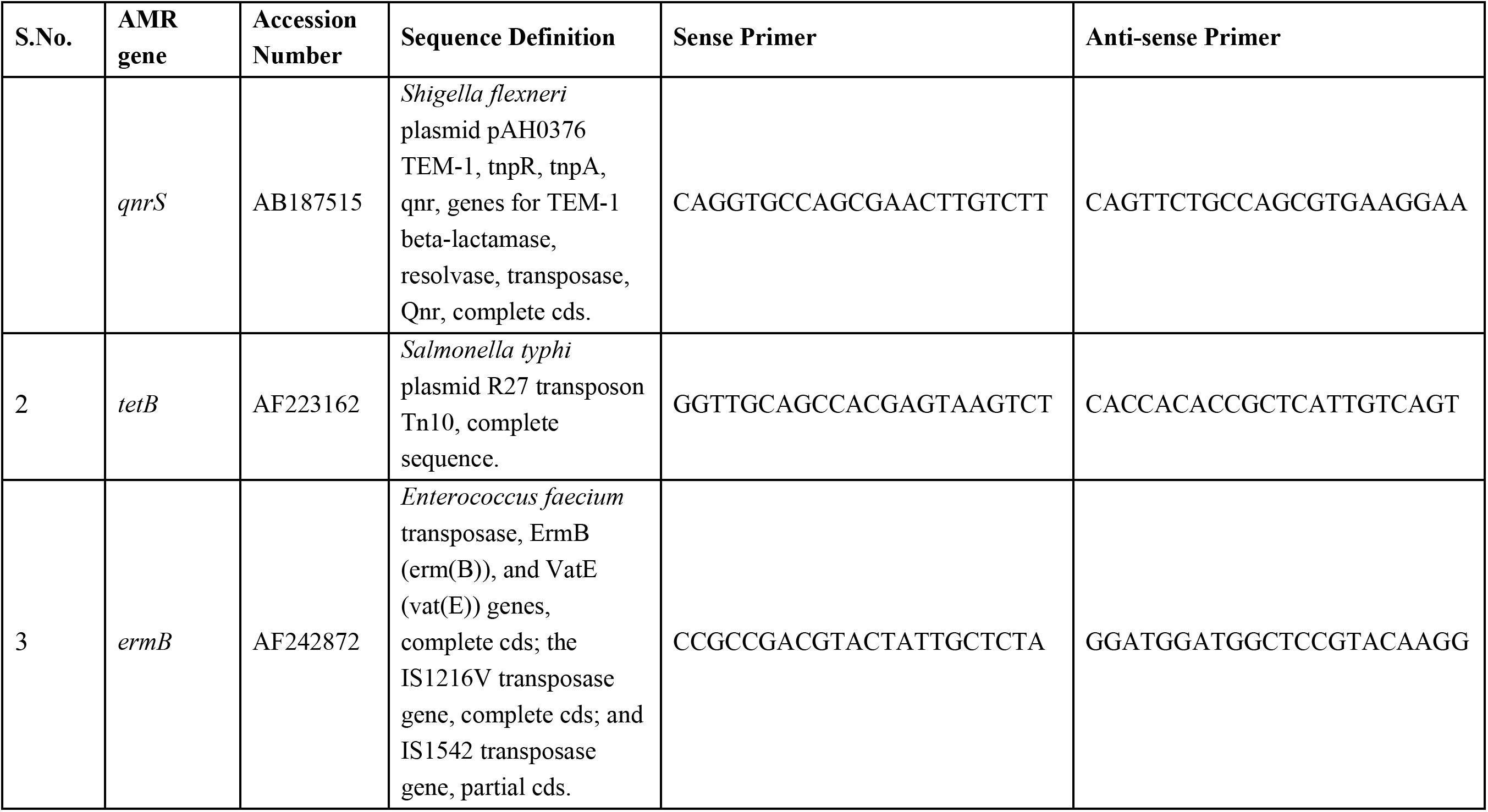
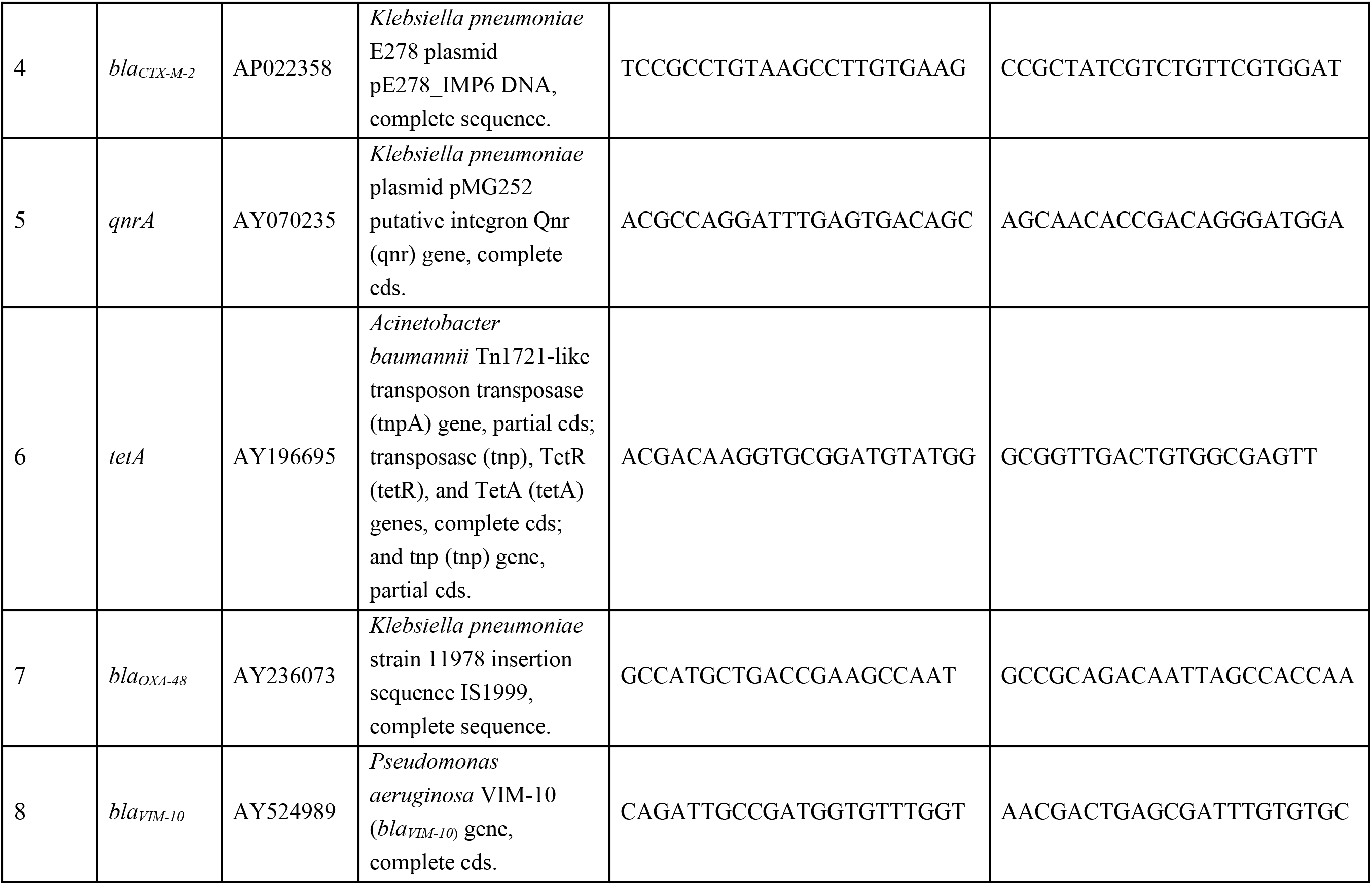
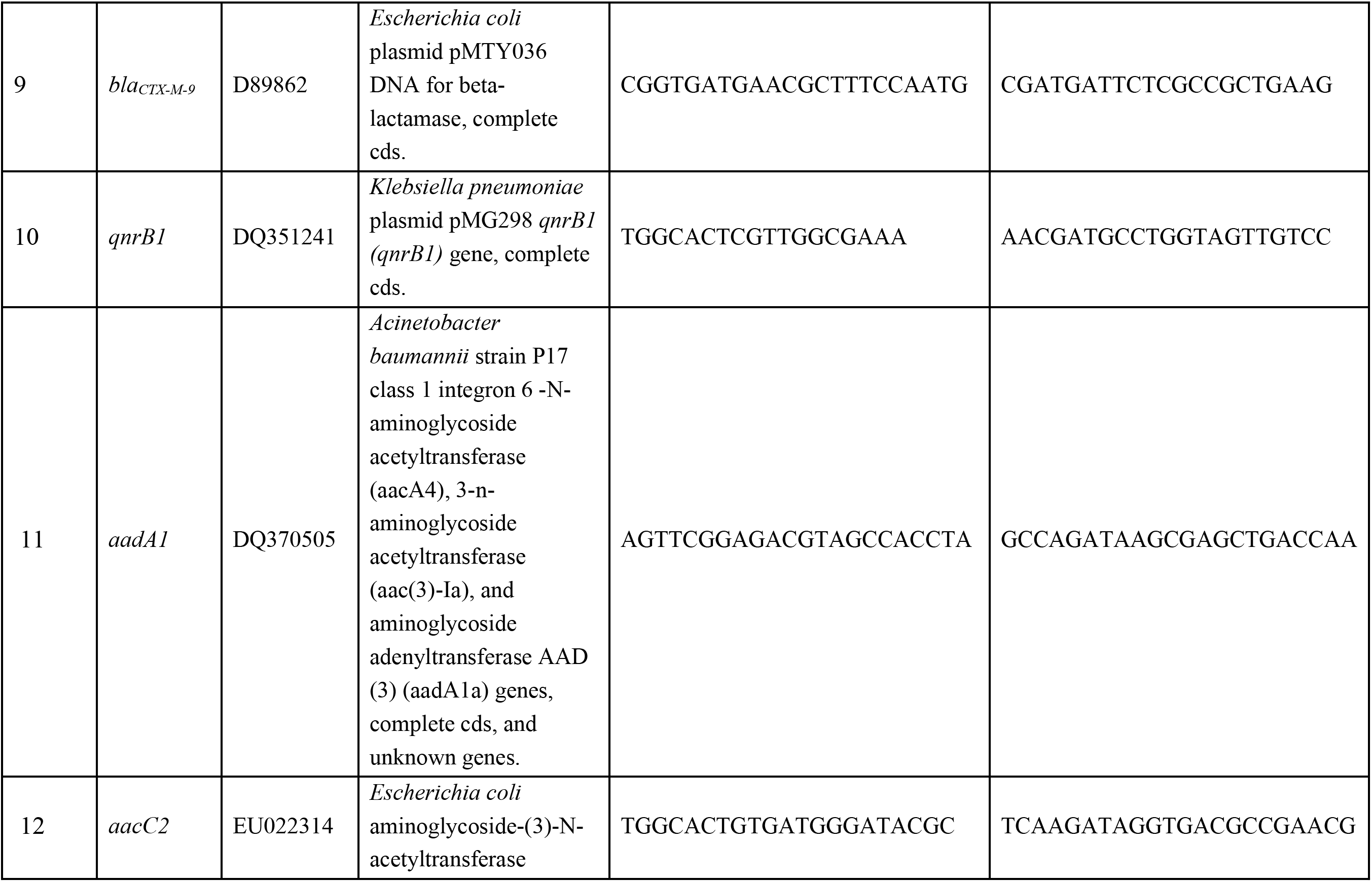
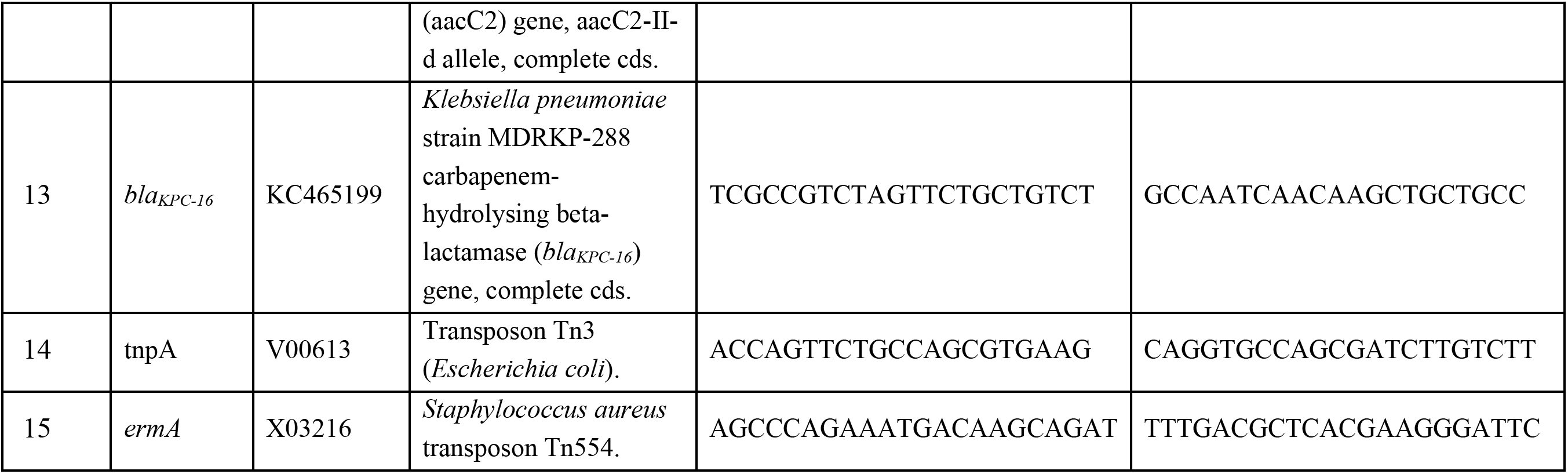
Primer sets listed below were evaluated during assay development but were excluded from the final multiplex PCR panel because they failed to yield reproducible amplification or showed suboptimal performance under optimised single-tube multiplex conditions.

**Supplementary Table S5:** Primer concentrations used for multiplex PCR.

| **S.No.** | **Target** | **Target category** | **Final concentration (µM)** |
|---|---|---|---|
| 1 | *bla<sub>SHV/LEN/OKP</sub>* | Class A β-lactamase | 0.8 |
| 2 | *bla<sub>CMY-6</sub>* | AmpC β-lactamase (Class C) | 0.8 |
| 3 | *bla<sub>NDM-1</sub>* | Carbapenemase (Class B metallo-β-lactamase) | 0.4 |
| 4 | *bla<sub>CTX-M-1</sub>* | CTX-M-1 group extended-spectrum β-lactamase | 0.4 |
| 5 | *bla<sub>TEM</sub>* | β-lactamase (Class A) | 0.4 |
| 6 | *qnrB1* | Plasmid-mediated quinolone resistance (PMQR) | 0.4 |
| 7 | *bla<sub>OXA-48</sub>* | Carbapenemase (Class D β-lactamase) | 0.4 |
| 8 | *silP* | Heavy-metal resistance determinant | 0.4 |
| 9 | 16S rRNA | Universal bacterial marker | 2.0 |
| 10 | 16S–ITS–23S rRNA operon (*rrn*) | Universal ribosomal operon marker | 2.0 |

\*Total volume of reaction: 50 µL

**Supplementary Table S6:** Predicted *in silico* coverage of the panel primers. Variants of each AMR gene family matched by the corresponding primer pair.

| **AMR gene** | **Cross-reacting variants** |
|---|---|
| *bla<sub>CMY</sub>* | 2, 4, 5, 6, 7, 12, 13, 14, 15, 16, 17, 18, 21, 22, 23, 24, 25, 26, 27, 28, 29, 30, 31, 32, 33, 34, 35, 36, 37, 38, 39, 40, 41, 42, 43, 44, 45, 46, 47, 48, 49, 50, 51, 53, 54, 55, 56, 57, 58, 59, 60, 61, 62, 63, 64, 65, 66, 67, 68, 69, 72, 73, 75, 76, 77, 78, 79, 80, 81, 84, 85, 86, 87, 90, 94, 95, 96, 97, 99, 102, 103, 104, 105, 106, 107, 108, 110, 111, 112, 113, 114, 116, 117, 118, 119, 121, 124, 125, 128, 129, 130, 131, 132, 133, 134, 136, 138, 139, 140, 141, 142, 143, 145, 146, 147, 148, 149, 150, 151, 152, 153, 154, 155, 156, 158, 160, 161, 162, 163, 164, 165, 166, 167, 168, 169, 170, 171, 173, 174, 185 |
| *bla<sub>CTX-M</sub>* | 1, 3, 11, 12, 22, 23, 30, 36, 42, 52, 54, 58, 60, 61, 62, 72, 80, 116, 136, 138, 146, 158, 162, 166, 167, 175, 187, 203, 204, 211, 220, 222, 236 |
| *bla<sub>NDM</sub>* | 1, 2, 3, 4, 5, 6, 7, 8, 9, 10, 11, 13, 14, 15, 16a, 16b, 17, 18, 19, 20, 21, 22, 23, 24, 25, 27, 28, 29, 33, 34, 35, 36, 37, 38, 39, 40 |
| *bla<sub>OXA</sub>* | 48, 162, 163, 181, 199, 204, 232, 244, 245, 247, 252, 370, 405, 416, 438, 439, 484, 505, 514, 515, 517, 519, 538, 546, 547, 566, 567, 731, 788, 793, 833, 894, 918, 920, 922, 923, 924, 929, 1038, 1039 |
| *bla<sub>SHV/LEN/OKP</sub>* | *SHV* 7, 14, 18, 27, 29, 30, 33, 34, 35, 36, 38, 40, 45, 51, 52, 56, 62, 63, 71, 72, 73, 77, 79, 80, 81, 82, 83, 85, 92, 93, 94, 96, 99, 103, 105, 108, 110, 111, 122, 132, 140, 141, 144, 150, 151, 157, 162, 163, 167, 168, 178, 187, 188, 189, 190, 191, 193, 195, 196, 198, 202, 204, 206, 207, 209, 213, 216, 219, 222, 224 |
|  | *LEN* 1, 2, 3, 4, 5, 6, 7, 8, 9, 10, 11, 12, 13, 14, 15, 16, 17, 18, 19, 20, 21, 22, 23, 24, 25, 26, 27, 28, 30, 31, 32, 34, 35, 36, 37, 38, 39, 40, 41, 42, 51, 55, 58 |
|  | *OKP-B* 1, 2, 3, 4, 5, 6, 7, 8, 9, 10, 12, 13, 14, 16, 17, 18, 19, 20, 21, 22, 23, 34, 40, 41, 45 |
| *bla<sub>TEM</sub>* | 1, 4, 7, 11, 12, 19, 40, 53, 54, 59, 60, 61, 63, 67, 68, 71, 72, 73, 75, 78, 80, 81, 82, 83, 84, 85, 86, 87, 89, 90, 91, 92, 93, 94, 96, 101, 109, 110, 111, 112, 113, 114, 115, 117, 118, 120, 121, 123, 125, 129, 130, 131, 133, 134, 137, 138, 139, 142, 146, 149, 150, 151, 152, 153, 154, 155, 156, 157, 159, 160, 162, 167, 168, 171, 177, 181, 183, 184, 185, 187, 188, 189, 190, 191, 195, 197, 199, 205, 211, 213, 219, 244 |
| *qnrB* | 1, 3, 6, 13, 14, 15, 16, 17, 18, 20, 23, 26, 41, 57, 58, 66, 74, 75, 76, 77 |
| *silP* | *silP* |

**Supplementary Table S7:** Summary of sequencing runs and replicates.

| **Sample type** | **Number of replicates** | **Sequencing platform** | **Flowcell type used in nanopore sequencing** | **Library Preparation Method** | **Input DNA** | **Number of reads** |
|---|---|---|---|---|---|---|
| Clinical isolates | 2 | Nanopore | MinION R10.4/<br>flongle R10.4 | Ligation Kit | 50 ng | BR1: 169275<br>BR2: 176732 |
| Zymo Microbial DNA | 2 | Nanopore | Minion/<br>flongle R10.4 | Ligation Kit | 50 ng | BR1: 112810<br>BR2: 22783 |
| Spiked blood | 2 | Nanopore | MinION R10.4/<br>flongle R10.4 | Ligation Kit | 50 ng | 10<sup>6</sup> CFU/mL: (BR1: 409938, BR2: 42353)<br>10<sup>5</sup> CFU/mL: (BR1: 21440, BR2: 1393)<br>10<sup>3</sup> CFU/mL: (BR1: 65411, BR2: 4211)<br>10<sup>2</sup> CFU/mL: (BR1: 176245, BR2: 603543)<br>10<sup>1</sup> CFU/mL: (BR1: 14720, BR2: 2713) |
| Environmental Samples | 2 or 3 | Nanopore | MinION R10.4/<br>Flongle R10.4 | Ligation Kit | 50 ng | See **Supplementary Table S9** for replicate-specific sequencing statistics |

**BR:** Biological replicate

**Supplementary Table S8:**
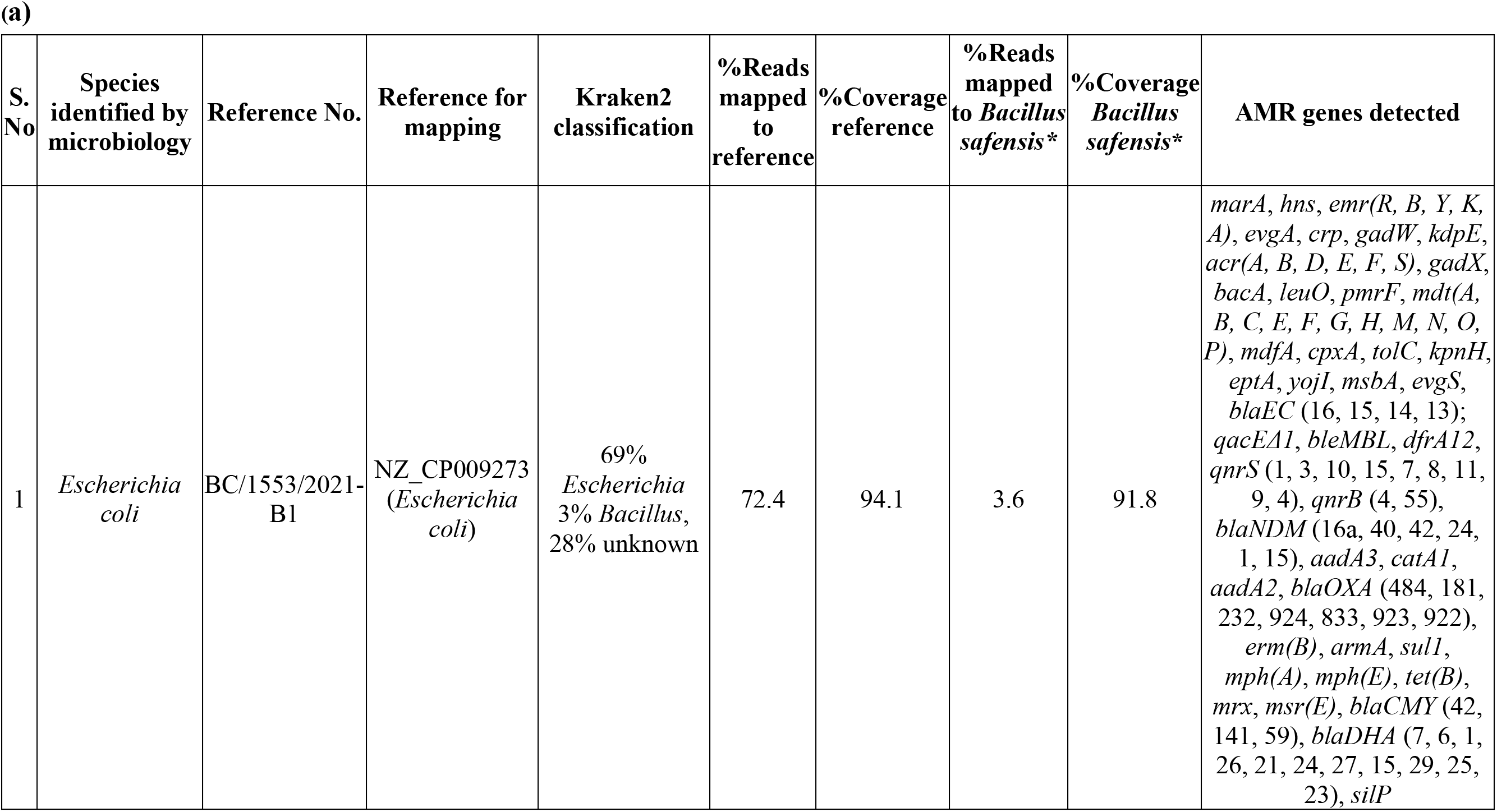
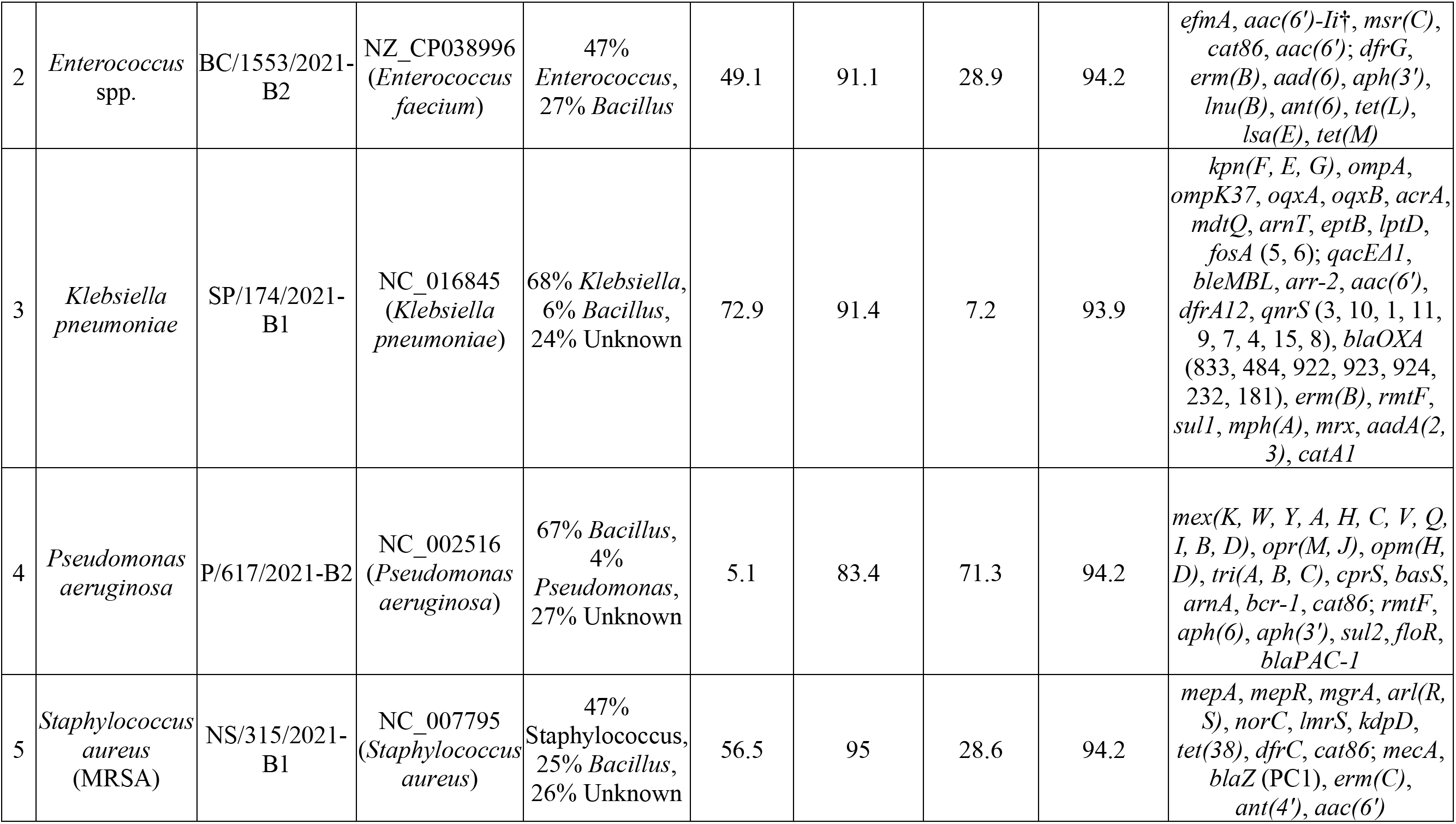
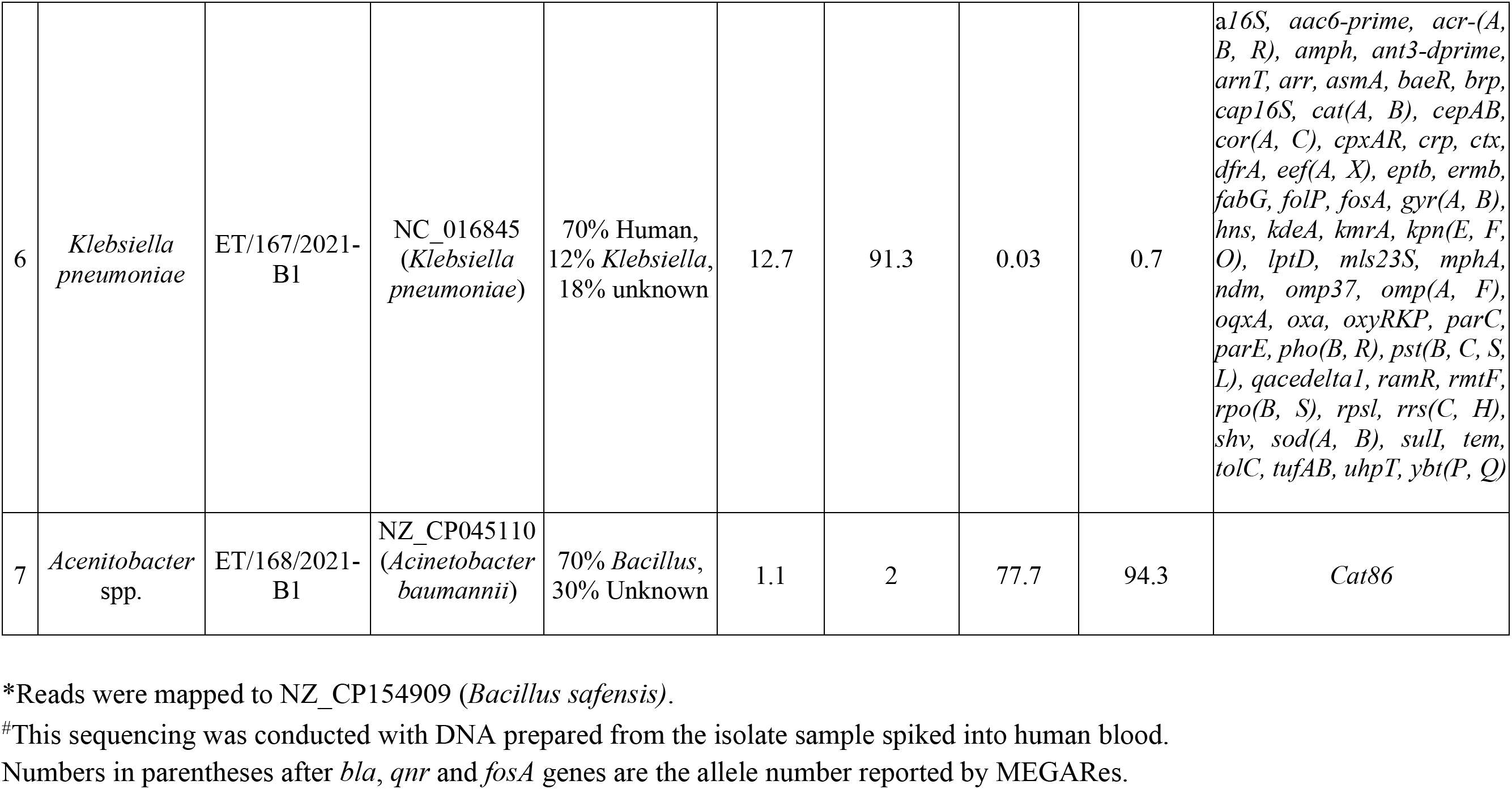
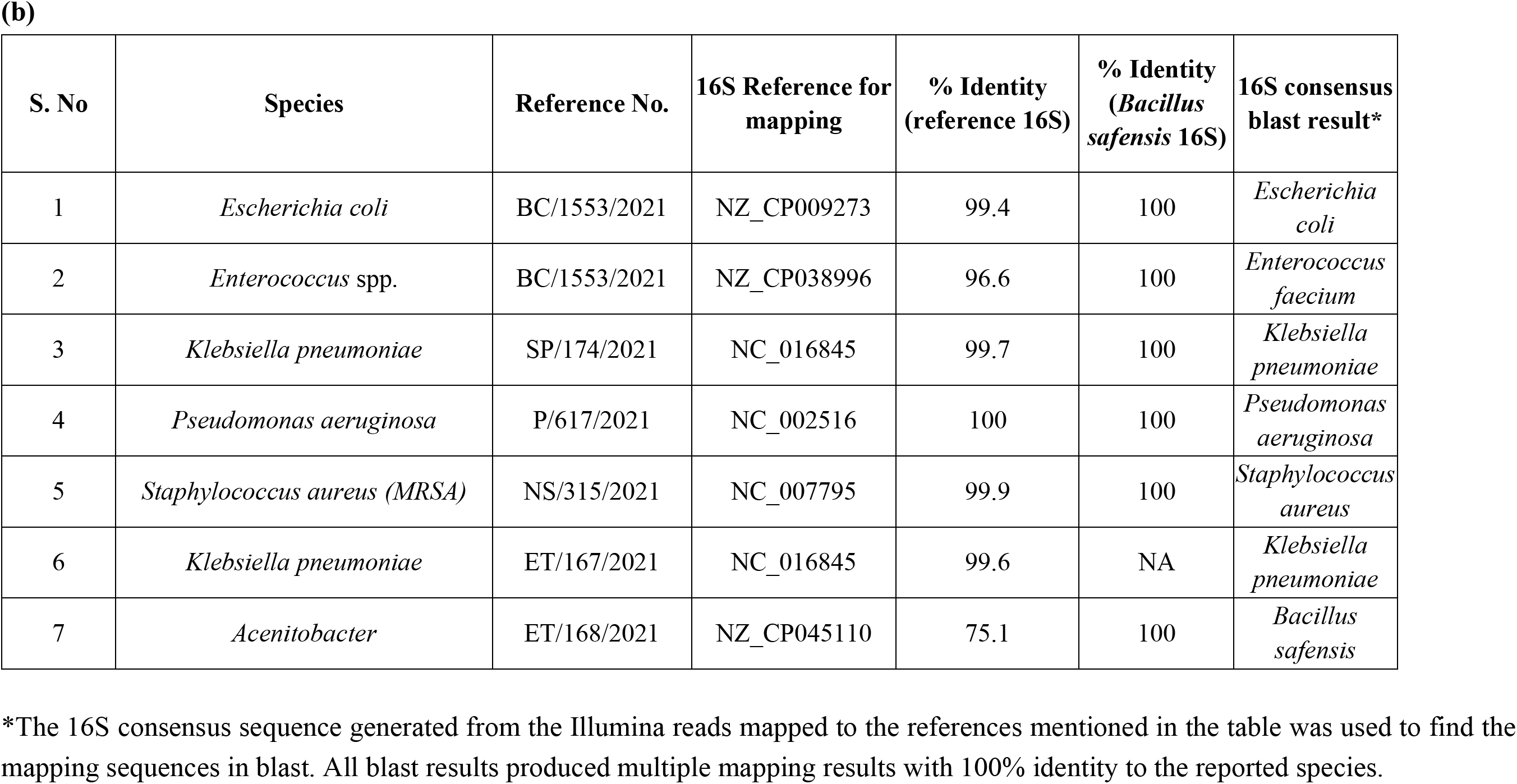
**(a)** Bacterial species and AMR gene detection results from Illumina whole-genome sequencing of the clinical isolates. **(b)** Illumina reads mapped to the 16S rRNA reference genes

| S. No | Species identified by microbiology | Reference No. | Reference for mapping | Kraken2 classification | %Reads mapped to reference | %Coverage reference | %Reads mapped to *Bacillus safensis\** | %Coverage *Bacillus safensis\** | AMR genes detected |
|---|---|---|---|---|---|---|---|---|---|
| 1 | *Escherichia coli* | BC/1553/2021-B1 | NZ_CP009273 (*Escherichia coli*) | 69% *Escherichia* 3% *Bacillus*, 28% unknown | 72.4 | 94.1 | 3.6 | 91.8 | *marA*, *hns*, *emr(R, B, Y, K, A)*, *evgA*, *crp*, *gadW*, *kdpE*, *acr(A, B, D, E, F, S)*, *gadX*, *bacA*, *leuO*, *pmrF*, *mdt(A, B, C, E, F, G, H, M, N, O, P)*, *mdfA*, *cpxA*, *tolC*, *kpnH*, *eptA*, *yojI*, *msbA*, *evgS*, *blaEC* (16, 15, 14, 13); *qacEΔ1*, *bleMBL*, *dfrA12*, *qnrS* (1, 3, 10, 15, 7, 8, 11, 9, 4), *qnrB* (4, 55), *blaNDM* (16a, 40, 42, 24, 1, 15), *aadA3*, *catA1*, *aadA2*, *blaOXA* (484, 181, 232, 924, 833, 923, 922), *erm(B)*, *armA*, *sul1*, *mph(A)*, *mph(E)*, *tet(B)*, *mrx*, *msr(E)*, *blaCMY* (42, 141, 59), *blaDHA* (7, 6, 1, 26, 21, 24, 27, 15, 29, 25, 23), *silP* |
| 2 | *Enterococcus* spp. | BC/1553/2021-B2 | NZ_CP038996 (*Enterococcus faecium*) | 47% *Enterococcus*, 27% *Bacillus* | 49.1 | 91.1 | 28.9 | 94.2 | *efmA*, *aac(6′)-Ii*†, *msr(C)*, *cat86*, *aac(6′)*; *dfrG*, *erm(B)*, *aad(6)*, *aph(3′)*, *lnu(B)*, *ant(6)*, *tet(L)*, *lsa(E)*, *tet(M)* |
| 3 | *Klebsiella pneumoniae* | SP/174/2021-B1 | NC_016845 (*Klebsiella pneumoniae*) | 68% *Klebsiella*, 6% *Bacillus*, 24% Unknown | 72.9 | 91.4 | 7.2 | 93.9 | *kpn(F, E, G)*, *ompA*, *ompK37*, *oqxA*, *oqxB*, *acrA*, *mdtQ*, *arnT*, *eptB*, *lptD*, *fosA* (5, 6); *qacEΔ1*, *bleMBL*, *arr-2*, *aac(6′)*, *dfrA12*, *qnrS* (3, 10, 1, 11, 9, 7, 4, 15, 8), *blaOXA* (833, 484, 922, 923, 924, 232, 181), *erm(B)*, *rmtF*, *sul1*, *mph(A)*, *mrx*, *aadA(2, 3)*, *catA1* |
| 4 | *Pseudomonas aeruginosa* | P/617/2021-B2 | NC_002516 (*Pseudomonas aeruginosa*) | 67% *Bacillus*, 4% *Pseudomonas*, 27% Unknown | 5.1 | 83.4 | 71.3 | 94.2 | *mex(K, W, Y, A, H, C, V, Q, I, B, D)*, *opr(M, J)*, *opm(H, D)*, *tri(A, B, C)*, *cprS*, *basS*, *arnA*, *bcr-1*, *cat86*; *rmtF*, *aph(6)*, *aph(3′)*, *sul2*, *floR*, *blaPAC-1* |
| 5 | *Staphylococcus aureus* (MRSA) | NS/315/2021-B1 | NC_007795 (*Staphylococcus aureus*) | 47% Staphylococcus, 25% *Bacillus*, 26% Unknown | 56.5 | 95 | 28.6 | 94.2 | *mepA*, *mepR*, *mgrA*, *arl(R, S)*, *norC*, *lmrS*, *kdpD*, *tet(38)*, *dfrC*, *cat86*; *mecA*, *blaZ* (PC1), *erm(C)*, *ant(4′)*, *aac(6′)* |
| 6 | *Klebsiella pneumoniae* | ET/167/2021-B1 | NC_016845 (*Klebsiella pneumoniae*) | 70% Human, 12% *Klebsiella*, 18% unknown | 12.7 | 91.3 | 0.03 | 0.7 | a*16S, aac6-prime, acr-(A, B, R), amph, ant3-dprime, arnT, arr, asmA, baeR, brp, cap16S, cat(A, B), cepAB, cor(A, C), cpxAR, crp, ctx, dfrA, eef(A, X), eptb, ermb, fabG, folP, fosA, gyr(A, B), hns, kdeA, kmrA, kpn(E, F, O), lptD, mls23S, mphA, ndm, omp37, omp(A, F), oqxA, oxa, oxyRKP, parC, parE, pho(B, R), pst(B, C, S, L), qacedelta1, ramR, rmtF, rpo(B, S), rpsl, rrs(C, H), shv, sod(A, B), sulI, tem, tolC, tufAB, uhpT, ybt(P, Q)* |
| 7 | *Acenitobacter* spp. | ET/168/2021-B1 | NZ_CP045110 (*Acinetobacter baumannii*) | 70% *Bacillus*, 30% Unknown | 1.1 | 2 | 77.7 | 94.3 | *Cat86* |
| **S. No** | **Species** | **Reference No.** | **16S Reference for mapping** | **% Identity (reference 16S)** | **% Identity (*Bacillus safensis* 16S)** | **16S consensus blast result\*** |  |  |  |
| 1 | *Escherichia coli* | BC/1553/2021 | NZ_CP009273 | 99.4 | 100 | *Escherichia coli* |  |  |  |
| 2 | *Enterococcus* spp. | BC/1553/2021 | NZ_CP038996 | 96.6 | 100 | *Enterococcus faecium* |  |  |  |
| 3 | *Klebsiella pneumoniae* | SP/174/2021 | NC_016845 | 99.7 | 100 | *Klebsiella pneumoniae* |  |  |  |
| 4 | *Pseudomonas aeruginosa* | P/617/2021 | NC_002516 | 100 | 100 | *Pseudomonas aeruginosa* |  |  |  |
| 5 | *Staphylococcus aureus (MRSA)* | NS/315/2021 | NC_007795 | 99.9 | 100 | *Staphylococcus aureus* |  |  |  |
| 6 | *Klebsiella pneumoniae* | ET/167/2021 | NC_016845 | 99.6 | NA | *Klebsiella pneumoniae* |  |  |  |
| 7 | *Acenitobacter* | ET/168/2021 | NZ_CP045110 | 75.1 | 100 | *Bacillus safensis* |  |  |  |

\*Reads were mapped to NZ_CP154909 (*Bacillus safensis)*.
<sup>#</sup>This sequencing was conducted with DNA prepared from the isolate sample spiked into human blood.
Numbers in parentheses after *bla*, *qnr* and *fosA* genes are the allele number reported by MEGARes.
\*The 16S consensus sequence generated from the Illumina reads mapped to the references mentioned in the table was used to find the mapping sequences in blast. All blast results produced multiple mapping results with 100% identity to the reported species.

**Supplementary Table S9:** Summary of the percentage of long and short reads for each sequencing run.

| **Sample type** | **Repeats** | **Total reads** | **Number of short reads** | **% Short reads** | **Number of long reads** | **% Long reads** | **% excluded reads<sup>#</sup>** |
|---|---|---|---|---|---|---|---|
| Clinical isolate mix | BR1 | 169275 | 6494 | 3.8 | 146393 | 86.5 | 9.7 |
|  | BR2 | 176732 | 21891 | 12.4 | 130864 | 74.0 | 13.6 |
| Zymo microbial community | BR1 | 112810 | NA* |  |  |  |  |
|  | BR2 | 22783 | NA* |  |  |  |  |
| Spike blood 10<sup>6</sup> CFU/mL | BR1 | 409938 | 140500 | 34.3 | 96209 | 23.5 | 42.3 |
|  | BR2 | 42353 | 21622 | 51.1 | 12131 | 28.6 | 20.3 |
| Spike blood 10<sup>5</sup> CFU/mL | BR1 | 21440 | 8651 | 40.3 | 9220 | 43.0 | 16.6 |
|  | BR2 | 1393 | 602 | 43.2 | 543 | 39.0 | 17.8 |
| Spike blood 10<sup>3</sup> CFU/mL | BR1 | 65411 | 17865 | 27.3 | 30678 | 46.9 | 25.8 |
|  | BR2 | 4211 | 1228 | 29.2 | 1775 | 42.2 | 28.7 |
| Spike blood 10<sup>2</sup> CFU/mL | BR1 | 176245 | 33087 | 18.8 | 67115 | 38.1 | 43.1 |
|  | BR2 | 603543 | 442320 | 73.3 | 51288 | 8.5 | 18.2 |
| Spike blood 10<sup>1</sup> CFU/mL | BR1 | 14720 | 4930 | 33.5 | 3480 | 23.6 | 42.9 |
|  | BR2 | 2713 | 840 | 31.0 | 859 | 31.7 | 37.4 |
| WWS Raja Canal STP Influent | BR1<br>TR1 | 24982 | 5823 | 23.3 | 1462 | 5.9 | 70.8 |
|  | BR1<br>TR2 | 321411 | 72698 | 22.6 | 25821 | 8.0 | 69.3 |
|  | BR2<br>TR1 | 87616 | 19559 | 22.3 | 5411 | 6.2 | 71.5 |
|  | BR2<br>TR2 | 129242 | 57642 | 44.6 | 33039 | 25.6 | 29.8 |
| WWS Raja Canal STP Effluent | BR1<br>TR1 | 12347 | 980 | 7.9 | 94 | 0.8 | 91.3 |
|  | BR1<br>TR2 | 13274 | 4086 | 30.8 | 890 | 6.7 | 62.5 |
|  | BR2 | 328630 | 32187 | 9.8 | 5893 | 1.8 | 88.4 |
| WWS Bagalur Lake Inlet | BR1 | 2492 | 98 | 3.9 | 347 | 13.9 | 82.1 |
|  | BR2 | 1064776 | 318446 | 29.9 | 342221 | 32.1 | 38.0 |
| WWS Bagalur Lake Outlet | BR1 | 1593 | 71 | 4.5 | 1054 | 66.2 | 29.4 |
|  | BR2 | 14271 | 442 | 3.1 | 9430 | 66.1 | 30.8 |
|  | BR3 | 122930 | 442 | 0.4 | 38837 | 31.6 | 68.0 |
| WWS Bagalur open well | BR1 | 93468 | 32520 | 34.8 | 7234 | 7.7 | 57.5 |
|  | BR2 | 17621 | 4559 | 25.9 | 7620 | 43.2 | 30.9 |
|  | BR3 | 153343 | 2943 | 1.9 | 128414 | 83.7 | 14.3 |

**BR**, biological replicate derived from independent DNA extraction; **TR**, technical replicate generated from independent multiplex PCR reactions and sequencing runs using the same extracted DNA preparation. Technical replicates were used to assess workflow reproducibility and were not treated as independent biological observations in statistical analyses.
\*Read length-based filtering was not performed since most of the reads (>95%) were within long reads.
<sup>#</sup>Read lengths that do not belong to long reads (1250-1750) and short reads (500-1000 bp)

**Supplementary Table S10:** Wastewater sample composition. The numbers represent the average percentage of reads mapping to each genus.

| **16S genus** | **Raja Canal influent** | **Raja Canal effluent** | **Bagalur Lake Inlet** | **Bagalur Lake Outlet** | **Bagalur Open well** |
|---|---|---|---|---|---|
| Unclassified | 38.17 | 35.76 | 59.07 | 48.54 | 64.32 |
| *Acidovorax* | 2.25 | 0.45 | 0.88 |  | 0.42 |
| *Acinetobacter* |  |  | 4.83 | 1.57 |  |
| *Agriterribacter* |  |  |  |  | 0.50 |
| *Aliarcobacter* | 28.92 | 2.52 |  |  | 1.00 |
| *Aquabacterium* | 1.19 |  | 0.58 | 12.64 |  |
| *Arcobacter* | 6.54 | 1.28 |  |  |  |
| *Armatimonas* |  |  |  |  | 10.98 |
| *Calidifontimicrobium* |  |  |  | 0.44 |  |
| *Cloacibacterium* | 2.35 | 5.34 |  |  |  |
| *Comamonas* | 0.66 |  |  | 0.43 |  |
| *Curvibacter* |  | 2.85 | 1.19 | 0.83 |  |
| *Cyanobium* |  |  | 5.83 | 6.12 |  |
| *Dechloromonas* | 2.36 | 1.37 |  |  | 0.41 |
| *Flavobacterium* |  | 16.12 |  |  |  |
| *Fluviibacter* |  | 12.43 |  |  |  |
| *Fundidesulfovibrio* |  |  |  |  | 1.43 |
| *Hydrogenophaga* |  | 0.35 | 2.58 |  |  |
| *Limnohabitans* |  | 0.46 | 3.45 |  |  |
| *Macromonas* | 1.61 |  |  |  |  |
| *Methyloradius* |  | 0.35 |  |  |  |
| *Parasegetibacter* |  |  |  |  | 1.77 |
| *Polynucleobacter* |  | 1.67 |  |  | 1.11 |
| *Pseudomonas* | 0.27 | 3.39 | 1.19 | 0.48 |  |
| *Ramlibacter* |  |  | 1.18 | 0.33 |  |
| *Rhodoferax* |  |  | 0.66 | 1.20 |  |
| *Schlegelella* |  |  |  | 8.30 |  |
| *Sideroxyarcus* |  |  |  | 0.62 | 0.35 |
| *Synechococcus* |  |  | 2.92 | 3.50 |  |
| *Terrimonas* |  | 0.40 |  |  |  |
| *Thauera* | 1.69 |  |  |  |  |
| *Vogesella* |  |  |  | 0.37 |  |
| Other bacteria | 14.00 | 15.24 | 15.64 | 14.99 | 17.70 |

